# tRNA-derived fragments and microRNAs in the maternal-fetal interface of a mouse maternal-immune-activation autism model

**DOI:** 10.1101/2019.12.20.884650

**Authors:** Zhangli Su, Elizabeth L. Frost, Catherine R. Lammert, Roza K. Przanowska, John R. Lukens, Anindya Dutta

## Abstract

tRNA-derived small fragments (tRFs) and tRNA halves have emerging functions in different biological pathways, such as regulating gene expression, protein translation, retrotransposon activity, transgenerational epigenetic changes and response to environmental stress. However, small RNAs like tRFs and microRNAs in the maternal-fetal interface during gestation have not been studied extensively. Here we investigated the small RNA composition of mouse placenta/decidua, which represents the interface where the mother communicates with the fetus, to determine whether there are specific differences in tRFs and microRNAs during fetal development and in response to maternal immune activation (MIA). Global tRF expression pattern, just like microRNAs, can distinguish tissue types among placenta/decidua, fetal brain and fetal liver. In particular, 5’ tRNA halves from tRNA^Gly^, tRNA^Glu^, tRNA^Val^ and tRNA^Lys^ are abundantly expressed in the normal mouse placenta/decidua. Moreover, tRF and microRNA levels in the maternal-fetal-interface change dynamically over the course of embryonic development. To see if stress alters non-coding RNA expression at the maternal-fetal interface, we treated pregnant mice with a viral infection mimetic, which has been shown to promote autism-related phenotypes in the offspring. Acute changes in the levels of specific tRFs and microRNAs were observed 3-6 hours after MIA and are suppressed thereafter. A group of 5’ tRNA halves is down-regulated by MIA, whereas a group of 18-nucleotide tRF-3a is up-regulated. In conclusion, tRFs show tissue-specificity, developmental changes and acute response to environmental stress, opening the possibility of them having a role in the fetal response to MIA.

## Introduction

Increasing evidence suggests that parental and gestational environmental exposure can affect the health of future generations through epigenetic modifications. In particular, emerging data have linked epigenetic modification to neurodevelopmental/neuropsychiatric disorders including autism spectrum disorder (ASD), schizophrenia and bipolar disorders in humans [1-4]. ASD is a prevalent condition and cannot be completely explained by genetic factors [5, 6]. Among the many environmental risk factors for ASD, infection-associated MIA (maternal immune activation) during pregnancy poses a strong risk factor for ASD in subsequent generations [4, 7-9] and has been recapitulated in animal models. For example, MIA-induced ASD can be modeled in mice by treating pregnant mothers with the viral mimetic polyinosinic:polycytidylic acid (Poly(I:C)) [10-15]. Poly(I:C) treatment induces systemic inflammation during pregnancy and, as a result, offspring from Poly(I:C)-treated dams develop many of the defining features of ASD. The MIA-triggered ASD phenotypes were found to be regulated by maternal immune pathways during gestation [10-15], suggesting close interaction between mother and the fetus. Such interaction takes place at the maternal-fetal interface, where fetal-derived placenta is bathed in the materal-derived decidua [16]. Indeed, maternal cytokines activated by MIA can be found in placenta shortly after the stress [17]. However, it remains to be determined whether other molecules such as short non-coding RNAs act at the maternal-fetal interface in parallel or downstream of maternal cytokines to regulate fetal development.

MicroRNAs (miRs) and tRNA-derived fragments (tRFs) are important short non-coding RNAs that regulate gene expression and are regulated by differentiation, development and environmental factors. Environmental factors such as diet, stress, and inflammation can promote the alteration of miRs and tRFs, which can directly lead to phenotypic alterations in future progeny [18-22]. However, whether non-coding RNAs are altered in models of neurodevelopmental disease has not been studied in great detail to date. miRs have been the most well studied small RNAs and have been proposed to regulate more than 50% of the transcriptome [23] including genes that regulate immune responses and neurodevelopment [24, 25]. On the other hand, tRFs are a newly identified class of non-coding RNAs that are beginning to be linked to various biological functions, including the regulation of gene expression and epigenetics [19, 26-45]. A group of tRFs, tRNA halves, generated by cleavage at tRNA anticodon loop, can be induced by different forms of stress including viral infection and RNases in innate immune responses [46-55]. Yet how environmental risk factors that have been linked to ASD, such as MIA, affect miR and tRF expression during fetal development has not been extensively studied.

Here we first determine whether tRFs and microRNAs at the maternal-fetal interface are differentially expressed in different tissues and altered during embryonic development. Finally, we test the hypothesis that maternal immune activation triggers change in the small RNAs, particularly tRFs and microRNAs, at the maternal-fetal interface. Overall our results suggest that tRFs are altered in a manner similar to microRNAs by tissue differentiation, by embryonic development and in response to MIA, suggesting that they could play a role in regulating the maternal-fetal interaction.

## Results

### Abundant 5’ tRNA fragments in placenta/decidua

First we set out to determine the small RNA composition in the maternal-fetal interface by global small RNA-sequencing approach. We isolated RNA from placenta/decidua of pregnant wild-type C57BL/6 female mice at E13.5 (embryonic day 13.5) and compared it with fetal brain and fetal liver from the same time point (Fig. S1). There was a high level of small RNAs around 30-50 nucleotides in placenta/decidua and fetal liver (Fig. 1B). Small RNAs were profiled by standard NGS library protocol (Fig. 1A-C). The cDNA ligated with adaptors was size selected to include inserts of 15-50 nucleotides and exclude mature tRNAs (Fig. 1C). For each tissue, we have 6 samples including 3 male and 3 female fetal mice from matched litters. 50-70% of sequenced short RNAs were microRNAs, compared to 6-25% being nuclear-derived tRFs and only 1-2% being mitochondrial tRFs (Fig. 1D). Since the mitochondria tRNA fragments are at a much lower abundance than the genomic (nuclear-encoded) tRNA fragments, we focused on genomic tRNA fragments and refer to them as “tRNA fragments” throughout the paper, unless specified otherwise. Very little difference was observed in microRNAs and genomic tRFs between fetal sexes (Fig. 1D). For reasons not clear to us, mitochondrial tRFs are significantly higher (p = 0.005 for fetal liver, p = 0.03 for fetal brain, unpaired two-tail student’s t test, 3 samples each group) in female fetal brain and fetal liver compared to the male counterparts (littermates from the same mother) (Fig. 1D). Moreover, there are tissue specific differences in the relative abundance of nuclear derived tRFs, which varied from 5% in the fetal brain to 20-25% in the placenta (Fig. 1D). Indeed, principle component analysis (PCA) showed clear clustering by tissue types by microRNAs (Fig. 1E) or tRNA fragments (Fig. 1F), suggesting both microRNAs and tRFs are significantly different among these three tissue types, though the difference in tRFs seems smaller than that in microRNAs.

**Figure 1.**
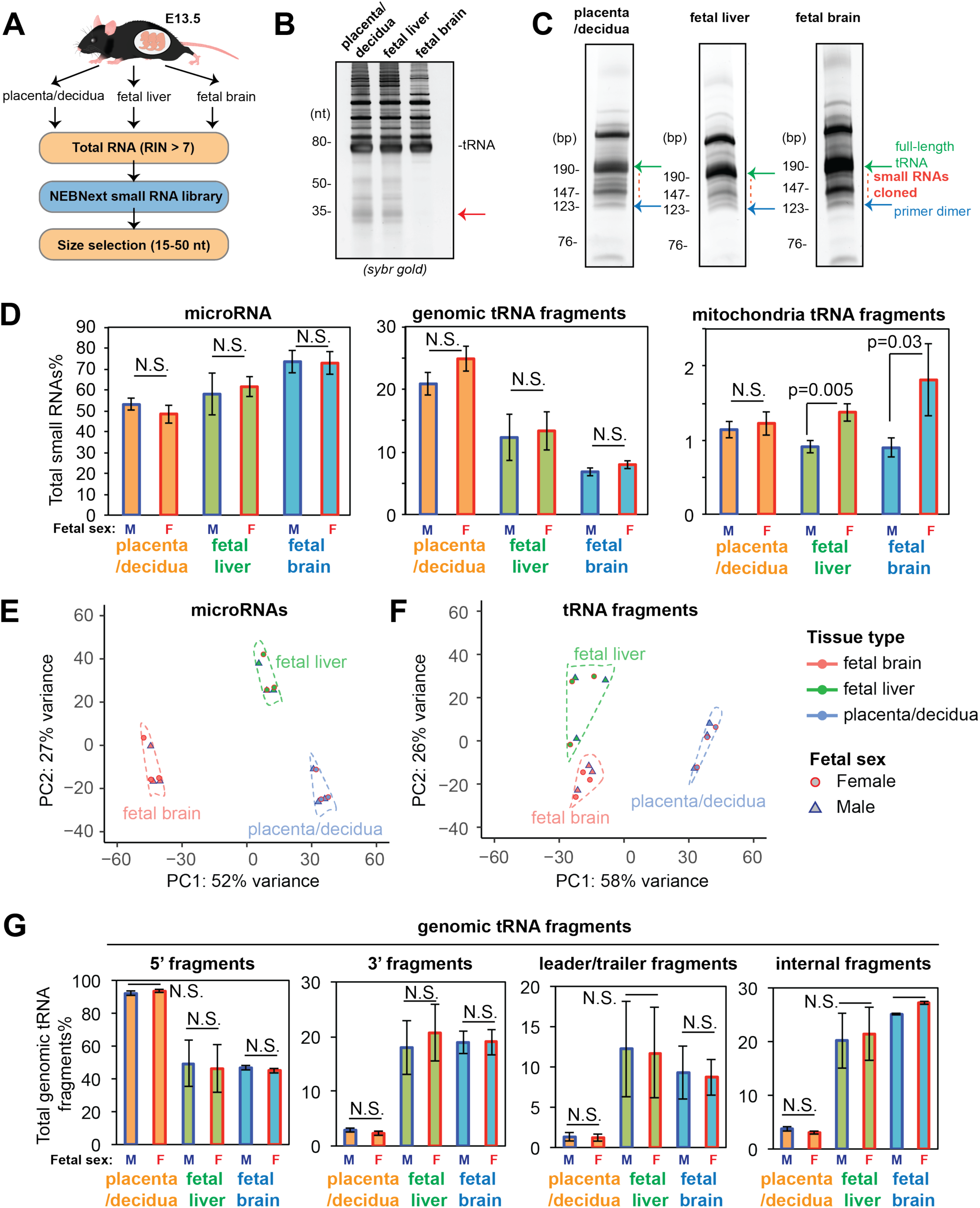
Small RNA profiling from three mouse tissues. (A) Brief workflow of sample collection and small RNA library preparation. Three tissue types (placenta/decidua, fetal liver and fetal brain) at embryonic day 13.5 (E13.5) were examined and divided by fetal sex (n = 3). (B) 5 μg total RNA from E13.5 tissues was resolved on 10% TBE-Urea-PAGE gel and stained with SybrGold. Red arrow indicates the additional small RNAs in placenta/decidua and fetal liver. (C) Representative SybrGold stained TBE-PAGE gel images of small RNA library. Red dashed line indicates the size selection for inserts between 15-50 nucleotide long RNAs. (D) Composition of different small RNAs (microRNAs, genomic tRFs and mitochondria tRFs). Y-axis shows percentage of each small RNA category in total mapped reads, error bars represent standard deviation. (E-F) Principle component analysis of microRNAs (E) and tRFs (F) from the three tissue types. (G) Composition of different tRF types (5’ fragments, 3’ fragments, leader/trailer fragments and internal fragments). Y-axis shows percentage of each tRF category in total tRF mapped reads, error bars represent standard deviation. P values are from unpaired two-tail student’s t test to compare sex differences in each tissue (N.S. = not significant, p value > 0.05).

We next determined whether the tRNA fragments seen in the fetus came from specific parts of the tRNAs. As described before [27, 56], tRFs can originate from the extreme 5’ end of mature tRNAs (tRF-5), the 3’ CCA end of mature tRNAs (tRF-3), the 3’ trailer sequence of the precursor tRNA (tRF-1), the 5’ leader sequence of the precursor tRNA or from an internal sequence of the mature tRNA (i-tRF). The placenta was particularly enriched in tRF-5 reads by small RNA-seq and dis-enriched in the other tRFs relative to the other tissues (Fig. 1G).

### 5’ tRNA halves in placenta/decidua from tRNA^Gly^, tRNA^Glu^, tRNA^Val^ and tRNA^Lys^

The high levels of tRNA 5’ fragments in the maternal-fetal interface are particularly interesting, given the previous reports about 5’ tRNA fragments involved in transgenerational transmission of epigenetic phenotypes in mice [19, 41, 45]. To further understand the nature of the short RNAs in placenta/decidua, we plotted the length distribution of different small RNAs. As expected, microRNA reads showed a sharp peak at 22 nucleotides in all three tissues (Fig. S2). In contrast to microRNAs, tRNA fragments in placenta/decidua show a major peak around 35 nucleotides (Fig. S2), which mostly come from the 5’ fragments (Fig. 2A). This is also consistent with the apparent RNA band around 35 nucleotides by SybrGold staining (Fig. 1B) and the high relative percentage of 5’ fragments by small RNA-seq in placenta/decidua (Fig. 1E). On the other hand, fragments from mitochondria tRNAs in placenta/decidua peaked at 21 and 41 nucleotide (Fig. S2), while fragments from snoRNA in placenta/decidua peaked at various sizes (Fig. S2). 5’ tRNA fragments from fetal liver also showed two major peaks at 35 and 32 nucleotide (Fig. S3A), but 5’ tRF from fetal brain displayed another major peak at 23 nucleotide (Fig. S3B). The 30-40 nucleotides long tRF-5s fall into the size range for 5’ tRNA halves, which is generated by cleavage at the anticodon loop. Thus 5’ tRNA halves constitute the major fraction of the tRF-5s in placenta and fetal liver, and are present in the brain, though smaller tRF-5s are quite notable in the last. Again no sex specific differences are obvious in the abundance of the tRFs.

**Figure 2.**
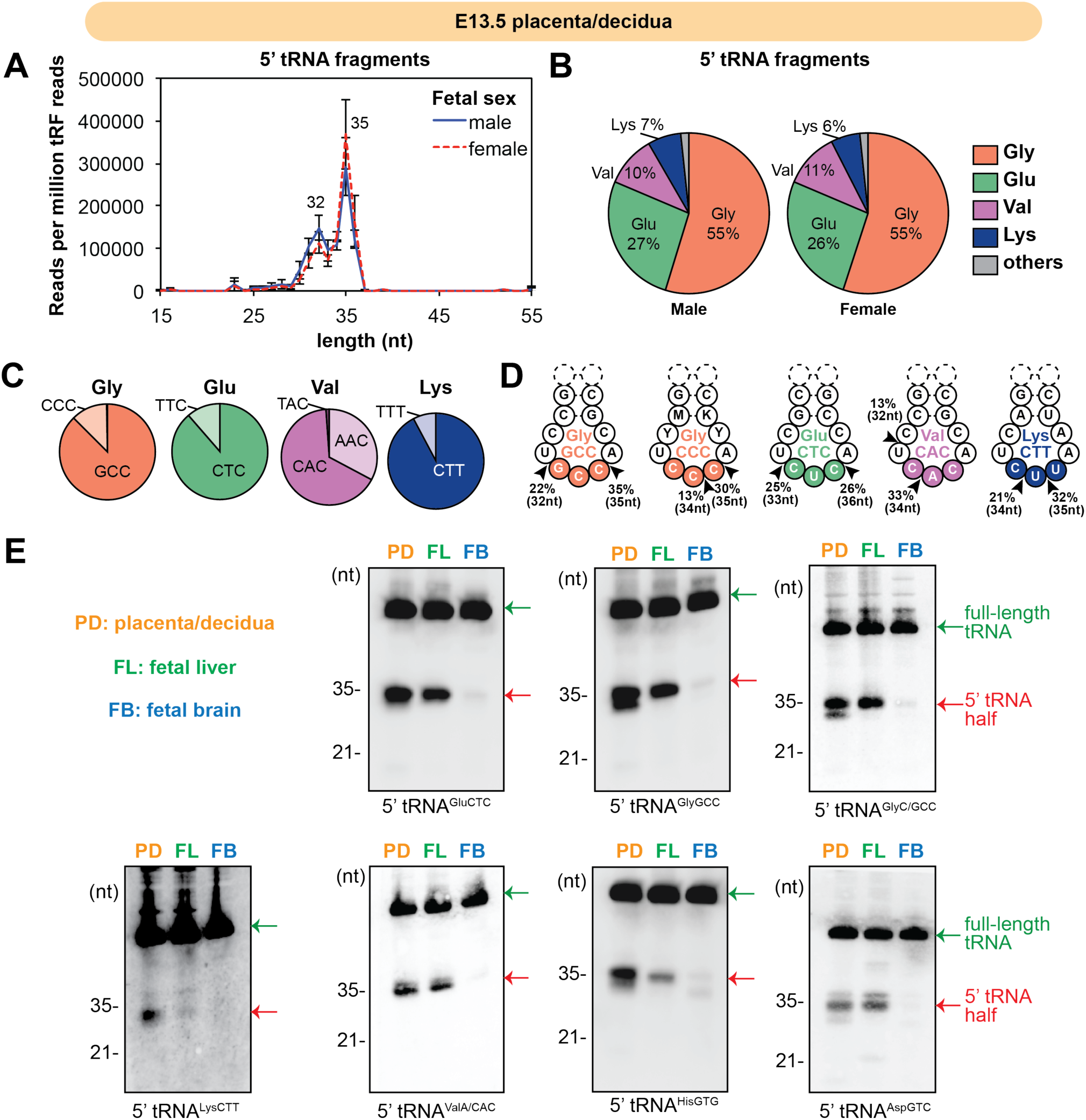
Abundant 5’ tRNA fragments in mouse maternal-fetal interface. (A-E) 5’ tRNA fragments in E13.5 placenta/decidua shows a high percentage of 5’ halves. (A) Length distribution of reads that map to 5’ end of tRNA (Y-axis: average reads per million total tRF reads). (B-C) Most 5’ tRNA fragments come from tRNA^Gly^, tRNA^Glu^, tRNA^Val^ and tRNA^Lys^. (C) The most representative anticodons were shown for each tRNA groups. (D) Arrows indicate the top two most abundant cleavage points, as indicated by the 3’ end position of 5’ tRNA reads. (E) Northern blots of 5’ tRNA halves in normal E13.5 tissues (PD: placenta/decidua, FL: fetal liver, FB: fetal brain). Arrows indicate full-length tRNAs (upper arrows) and 5’ tRNA halves of around 35 nucleotides (lower arrows).

The 5’ tRNA fragments (or halves) in placenta/decidua small RNA-seq reads come from four dominant tRNA groups (tRNA^Gly^, tRNA^Glu^, tRNA^Val^ and tRNA^Lys^), covering 99% of all 5’ tRNA fragments detected (Fig. 2B). More than 80% of 5’ halves of tRNA^Gly^ come from tRNA^GlyGCC^ and the rest come from tRNA^GlyCCC^ (Fig. 2C). Similarly, tRNA^GluCTC^, tRNA^ValCAC^ and tRNA^LysCTT^ give rise to the most abundant 5’ tRNA halves compared to the other anticodons for the same amino acid (Fig. 2C). Examination of the 3’ end position of these 5’ tRNA halves reveals most of them end in the anticodon loop, close to the anticodon triplet (Fig. 2D, arrows indicate the top two highest frequency of sequenced 3’ end). The other tRNAs that gave rise to tRNA halves include tRNA^Asp^, tRNA^His^, tRNA^Leu^ and tRNA^iMet^ (Fig. S3C). The very high abundance of 5’ tRNA halves in placenta/decidua and fetal liver is also confirmed by Northern blots (Fig. 2E). The abundance and selectivity of tRNA halves suggest undiscovered biology that might be associated with the high levels of 5’ tRNA halves in the maternal-fetal interface.

### 3’ tRNA fragments in placenta/decidua

3’ fragments from mature tRNA containing 3’ CCA sequence are another important groups of tRFs that have previously been shown to have various functions. The short 3’ fragments are also named tRF-3s, usually have length of 18 or 22 nucleotides and like microRNAs can repress genes by interacting with target mRNAs along with Argonaute and GW182 proteins [39, 57, 58]. They could also regulate protein translation [40] or inhibit retrotransposon activity [44]. The long 3’ fragments are also called 3’ tRNA halves, usually around 40 nucleotides, which have been less studied and shown to regulate apoptosis [59] and translation [60]. In particular, 3’ tRNA fragments from placenta/decidua have a peak around 39 nucleotides (Fig. 3A), which are 3’ tRNA halves derived from cleavage in the anticodon loop. These 3’ tRNA halves are not abundant in fetal liver and fetal brain (Fig. S3A and Fig. S3B). Northern blots confirmed the presence of 3’ tRNA halves from tRNA^Glu^ and tRNA^Asp^ in placenta/decidua (Fig. 3C), the two tRNA groups that have the highest read count of 3’ halves in small RNA-seq (Fig. 3B). In addition, the tRF-3s from all three tissues have two major peaks at 18 and 22 nucleotides (Fig. 3A, Fig. S3A and Fig. S3B), corresponding to the previously reported size of tRF-3a and tRF-3b [35, 39, 44, 57, 58]. The small tRF-3s (< 30 nucleotides) derived from tRNA^Ala^, tRNA^Val^, tRNA^Gln^, tRNA^Gly^, tRNA^Leu^, tRNA^Cys^, tRNA^Tyr^, tRNA^Thr^ and tRNA^Ser^ contribute to most of the tRF-3 reads (Fig. 3D and Fig. S5). To validate the presence of short tRF-3s, we developed a modified qRT-PCR method that ligates adaptor sequence to both ends of RNAs and detects small RNAs by RT-PCR primers that are unique to the adaptor-tRF junctional sequences (Fig. S4B). Modified qRT-PCR protocol successfully detects the 18 nucleotides long tRF-3a from tRNA^Ala^ and tRNA^Tyr^ (Fig. 3E), which fell below the detection limits for Northern blots (Fig. S4A). The same qRT-PCR method also detects microRNAs (mmu-miR-26a-5p and mmu-miR-21a-5p) and 5’ tRNA halves (Fig. 3E). Interestingly, 5’ tRNA half levels increased when the RNAs were first treated with T4 PNK (polynucleotide kinase) (Fig. 3E), while there is no similar change in miR or tRF-3 levels. This suggests the total level of 5’ tRNA halves is under-estimated by our small RNA-seq protocol due to a technical bias in detecting RNAs with terminal and internal modifications by small RNA-seq (further discussed in Discussion). A more detailed distribution of the different tRF types and parental tRNAs by small RNA-seq in each tissue type is shown in Fig. S5.

**Figure 3.**
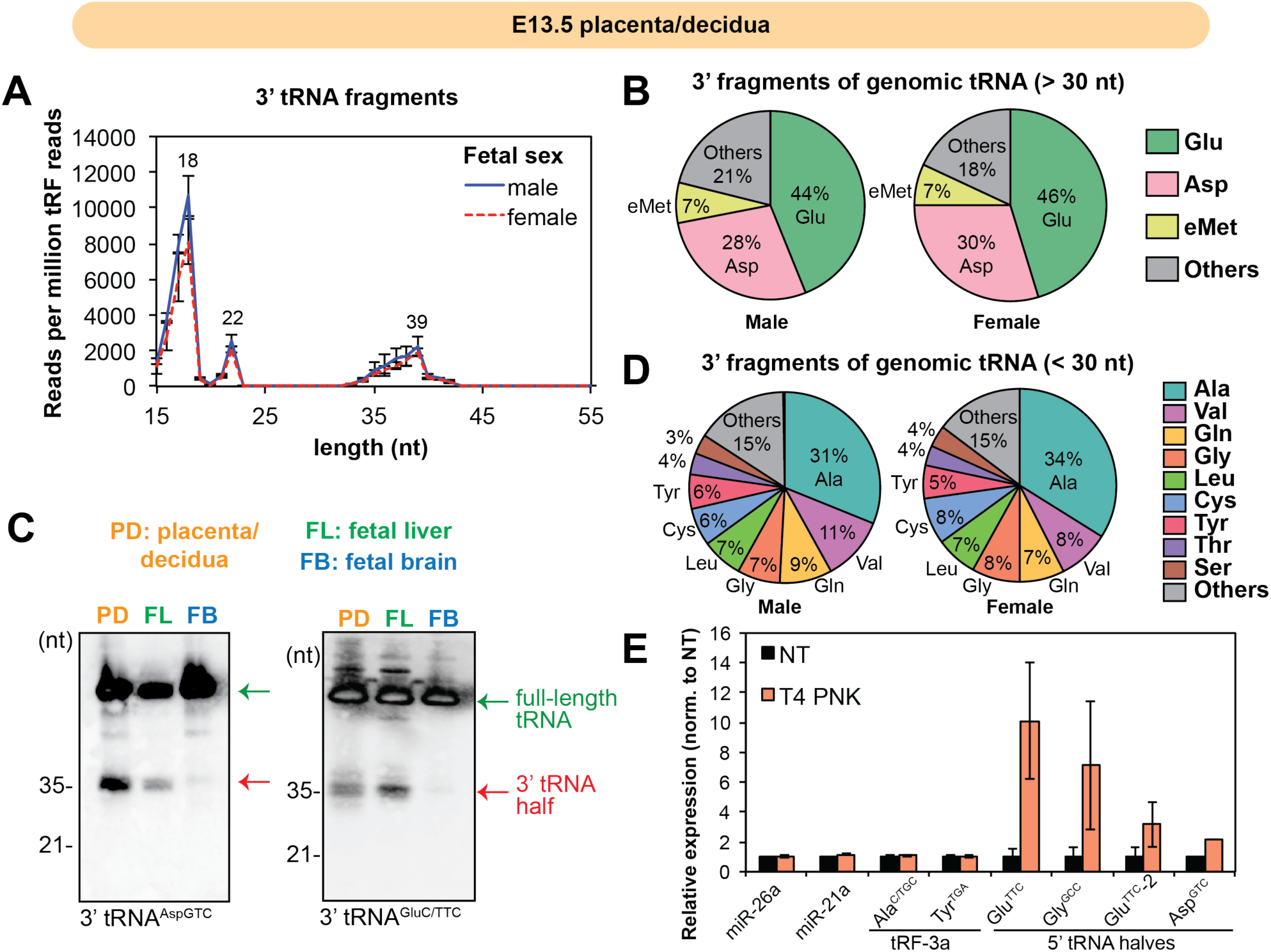
Detection of 3’ tRNA fragments in placenta/decidua. (A-C) 3’ tRNA fragments in E13.5 placenta/decidua shows distinct peaks by small RNA-seq for long (>30 nt) 3’ tRNA halves (B) and short (< 30 nt) tRF-3s (C). (A) Length distribution of reads that map to 3’ CCA end of tRNA (Y-axis: average reads per million total tRF reads. (B-C) Pie charts show percentage of parental tRNAs where long (> 30 nt) 3’ tRNA halves (B) and short (< 30 nt) tRF-3s (C) come from. (D) Northern blots of 3’ tRNA halves in normal E13.5 tissues (PD: placenta/decidua, FL: fetal liver, FB: fetal brain). Arrows indicate full-length tRNAs (upper arrows) and 3’ tRNA halves (lower arrows). (E) qRT-PCR detection of tRF-3 and 5’ tRNA halves in normal placenta/decidua sample (E12.5) after T4 PNK treatment to remove potential terminal modifications (NT: non treatment control). Error bars represent standard deviation from 3 biological replicates.

### Temporally dynamic expression of small RNAs in maternal-fetal interface

We next examined whether the repertoire of small RNAs expressed at the maternal-fetal interface changes over time from E12.5 to E18.5 (Fig. 4A). The lack of differential expression in short RNAs by fetal sex (Fig. S6) justifies pooling samples of both sexes for each time point. Principle component analysis shows that microRNAs and tRFs are relatively unchanged at E12.5, E13.5 and E14.5, but there is a substantial change at E18.5 (Fig. 4B). Thus like microRNAs, tRFs show significant change in expression at the maternal-fetal interface during fetal development. A heatmap of the dynamically expressed small RNAs identifies microRNAs and tRFs that are increasing or decreasing over time (Fig. 4C). The full list of dynamically expressed microRNAs and tRFs are in Supplementary Table 2. For example, mmu-miR-146b-5p decreased about 5-fold over the same period of time (Fig. 4D), and mmu-miR-215-5p increased about 10-fold from E12.5 to E18.5 (Fig. 4E).

**Figure 4.**
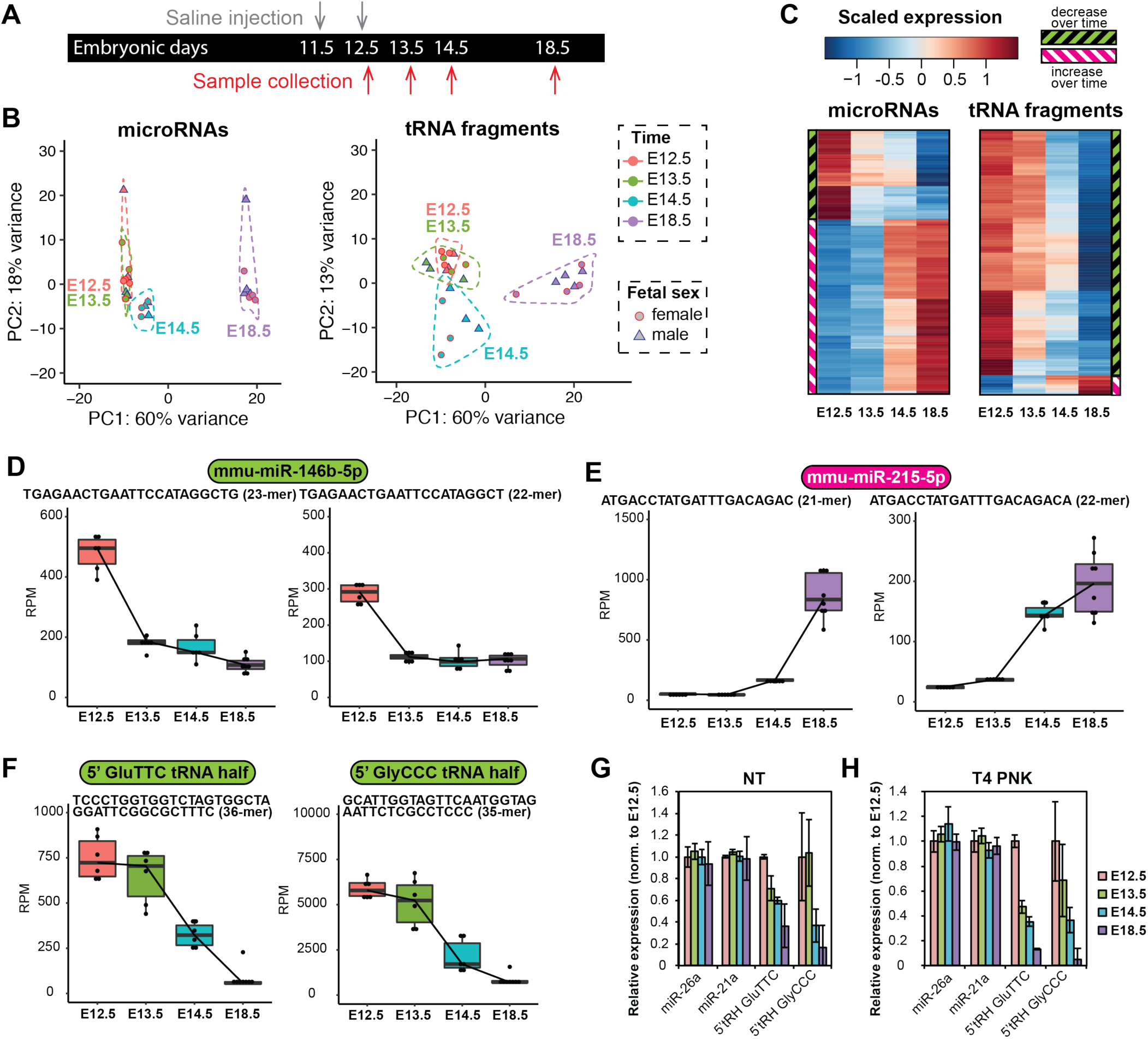
Dynamic expression of placental/decidual microRNAs and tRNA fragments. (A) Scheme of sample collection for this figure. Placenta/decidua control samples were collected at embryonic development days E12.5, E13.5, E14.5 and E18.5 (corresponding to the 3 hours, 24 hours, 48 hours, 144 hours time point control samples). (B) Principle component analysis of microRNAs and tRFs from placenta/decidua control samples at different time points by sequence-level analysis. (C) Heatmap shows dynamic expression of placental/decidual microRNAs and tRFs across embryonic development time. Each row in heatmap represents one unique sequence. Expression values were calculated by log10RPM (reads per million total mapped reads) and averaged from 6 samples at each time point and then scaled across row. For visualization purpose, only abundantly expressed miRs or tRFs (mean expression cut-off 100 RPM) that show time-dependent changes are shown in the heatmap. A complete list of dynamically expressed miRs and tRFs please refer to Table S2. (D-E) Examples of dynamically expressed microRNAs. mmu-miR-215-5p (D) is up-regulated over time and mmu-miR-146b-5p (E) is down-regulated over time. (F-H) Examples of dynamically expressed tRFs, including down-regulated 5’ halves from tRNA^GluTTC^ and tRNA^GlyCCC^. (D-F) Box plots showing RPM (reads per million total mapped reads) for specific miR or tRF sequence, with each dot represents one sample (n = 6 for each time point) and middle line represents median value. (G-H) qRT-PCR validation of temporal decrease of 5’ tRNA halves in both NT (no treatment control) and T4 PNK treatment samples. Error bars represent standard deviation from two biological replicates (male and female control samples pooled for each time point).

Among the temporally regulated placental/decidual tRFs, more of them are down-regulated than up-regulated over time (Fig. 4C) and most are 5’ tRNA halves (Supplementary Table 2). The down-regulated tRFs include abundant 5’ halves from tRNA^GluTTC^ and tRNA^GlyCCC^, which decreased 5-10 fold from E12.5 to E18.5 (Fig. 4F). The temporal decrease of 5’ tRNA halves was further validated by the modified qRT-PCR approach (Fig. 4G-H), and the trend was not affected by T4 PNK treatment. On the other hand, 5’ half from tRNA^HisGTG^ increased over this same time (Fig. S7), suggesting this is not a size selection artifact. Both 5’ unmodified and 5’ terminal guanosine-modified (G_-1_) tRNA^HisGTG^ halves show consistent up-regulation over time (Fig. S7). Overall, the developmental changes of placental/decidual small RNAs suggest that changes in the small RNAs could affect the interaction between the fetus and the mother.

### Changes in microRNAs and tRFs at the maternal-fetal interface in response to maternal immune activation: overall experimental design

Having observed the robust and dynamic expression of microRNAs and tRNA fragments at the maternal-fetal interface during normal gestation, we were interested to investigate whether these small RNAs could respond to maternal immune activation, which has been shown to increase the risk of neurodevelopmental disease in the offspring. In particular, MIA has been shown to increase the risk of autism and schizophrenia [4, 7-9]. To trigger systematic maternal immune activation during pregnancy, we treated pregnant mice with poly(I:C), a viral RNA mimetic, at E11.5 and E12.5, as previously described [11]. Importantly, this poly(I:C) treatment regimen promotes the development of autism-related phenotypes in the MIA-offspring [10-15]. Maternal-fetal interface (placenta/decidua) samples were collected at various time points (3, 6, 12, 24, 48, 144 hours) after the second injection at E12.5 (Fig. 5A). As a control, pregnant mice were injected with saline. This experimental set-up allows us to examine whether MIA triggers any changes in small RNA expression over time.

**Figure 5.**
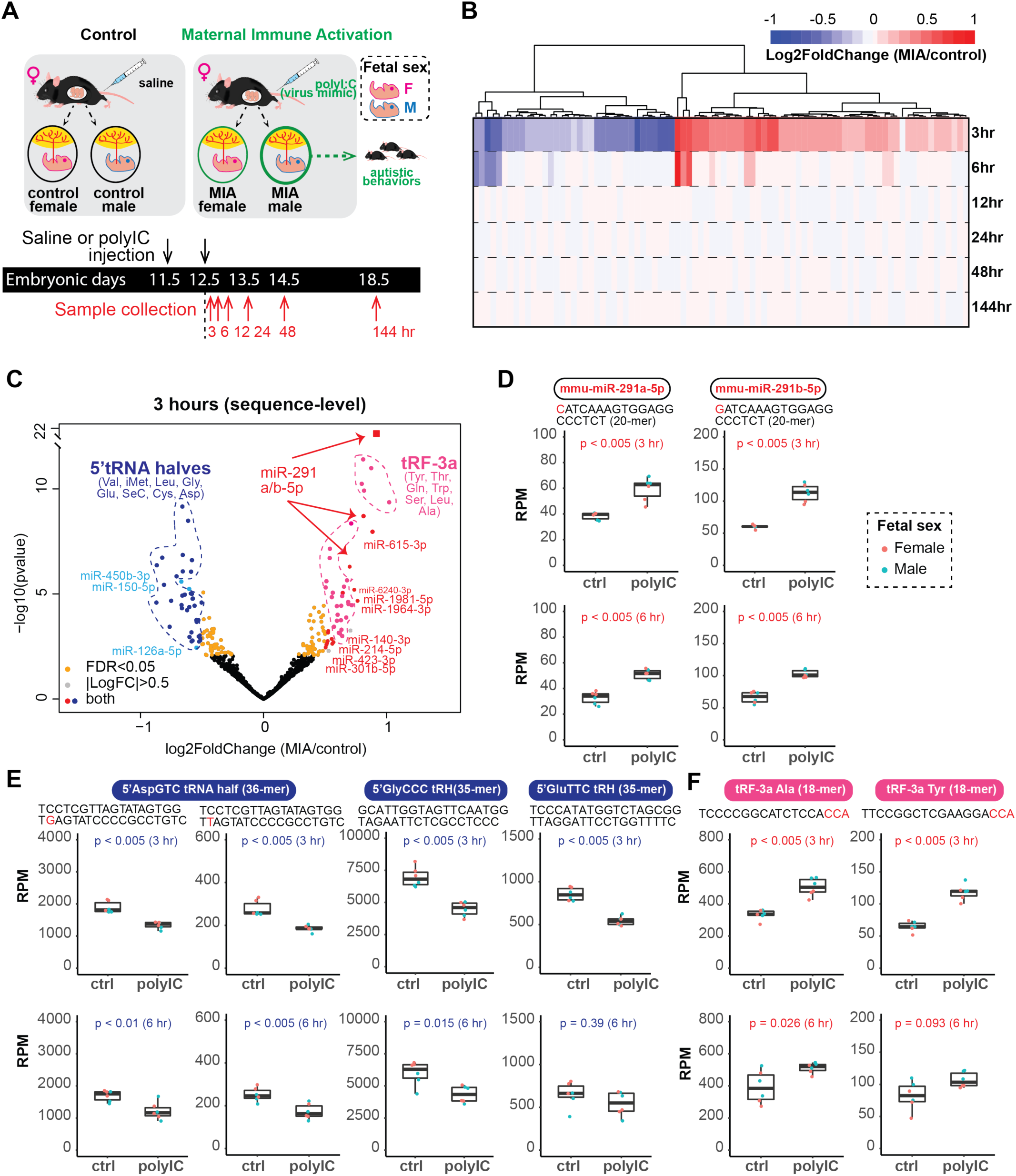
MIA elicits acute and transient changes in miR and tRF expression at the maternal-fetal interface. (A) Scheme of maternal immune activation (MIA) treatment and sample collection. MIA was triggered by Poly(I:C) injection of pregnant mice at E11.5 and E12.5, saline injection was used as a control. Placenta/decidua samples were collected 3 hours, 6 hours, 12 hours, 24 hours, 48 hours, 144 hours post second injection at E12.5. Fetal sex was determined by PCR. (B) Heatmap highlights early response to MIA stress in small RNA expression. Each column represents one microRNA or tRF. For visualization purpose, only miRs/tRFs that show significant MIA-dependent changes at any given time point (calculated by DESeq2, adjusted p < 0.05) are shown. Color indicates Log2 fold change (MIA over control) from DESeq2 estimation (apeglm shrinkage, fetal sex combined, n = 4 for 12 hours time point, n = 6 for all the other time points). (C) Volcano plot showing the MIA-dependent changes at the 3 hour time point. Significant miRs and tRF sequences (adjusted p < 0.05) are labeled. X-axis: Log2 fold change (MIA/control) from DESeq2 estimation (apeglm shrinkage, n = 6, fetal sex combined); Y-axis: -log10(p value). Each dot represents a unique sequence (with expression cut-off of 10 DESeq2 normalized reads per sample). Pink and blue circles highlight the tRF-3a and 5’ tRNA halves that are significantly altered by MIA. Full list of MIA-responsive miRs and tRFs refer to Table S3. (D-F) Box plots showing RPM (reads per million total mapped reads) for specific miR or tRF sequence, with each dot represents one sample (n = 6 for each time point, fetal sex is specified by dot color) and middle line represents median value. To compare the difference between control group and poly(I:C) (MIA) group, unpaired two-samples Wilcoxon test was performed (n = 6 for each group).

### Placental/decidual microRNAs and tRFs show early response to maternal immune activation

In order to find the placental/decidual small RNAs regulated by poly(I:C)-triggered maternal immune activation, differential analysis was performed for each time point between the poly(I:C) treatment group and the control group (male and female samples were combined to increase sample size). We found that small RNAs in the placenta/decidua displayed poly(I:C)-dependent changes as early as 3 hours post second poly(I:C) injection at E12.5 (Fig. S8A). Such changes detected at placenta/decidua diminished over time - there is less difference at 6 hours compared to 3 hours (Fig. S8B); at 12 hours and later time points, there is very little differentially expressed small RNAs between control and poly(I:C) treated groups (Fig. S8C-E), suggesting MIA very transiently alters small RNA expression at the maternal-fetal interface.

We focused on the 3 hour time point as this is the time point with the most significant MIA-dependent changes. At the 3 hour time point, we identified a group of placenta small RNAs (both microRNAs and tRFs) that are significantly altered by poly(I:C) treatment of pregnant mice (Fig. S9A). The MIA-induced change in this group of small RNAs gradually diminished over time (Fig. 5B, color indicates log2 fold change for each time point). Several tRF-3s (<30 nt) are up-regulated by MIA and some 5’ tRNA halves are down-regulated at 3 hours (Fig. S9A). Since tRFs of different lengths and classes are expected to have different functions, we further performed differential analysis at the individual sequence level (Fig. 5C, full list see Table S3). The most significantly altered microRNA at 3 hours is miR-291a/b-5p, which is up-regulated by MIA (Fig. 5C and Fig. S9A). miR-291a/b-5p still remain significantly up-regulated by MIA at 6 hour time point (Fig. 5D), but the difference disappears at 12 hours onwards. Furthermore, 5’ halves from tRNA^Asp^, tRNA^Gly^, tRNA^Glu^, tRNA^Val^, tRNA^iMet^, tRNA^Leu^ tRNA^SeC^ and tRNA^Cys^ are down-regulated by MIA at 3 hours (Fig. 5C and Fig. 5E). In particular, 5’ halves from tRNA^Asp^ continue to be significantly down-regulated at 6 hours (Fig. 5E). 5’ halves from tRNA^Gly^ and tRNA^Glu^ are also down-regulated, albeit the degree of down-regulation has diminished at 6 hours (Fig. 5E). On the other hand, 5’ half from tRNA^His^ is not decreased by MIA, but shows a slight increase at 3 hours (Fig. S9B). Finally, multiple 18-nucleotide long tRF-3a (including tRF-3^Tyr^, tRF-3^Gln^, tRF-3^Thr^, tRF-3^Leu^, tRF-3^Ser^, tRF-3^Trp^ and tRF-3^Ala^) are significantly up-regulated by MIA (Fig. 5C and Fig. 5F) but this diminishes by 6 hours (Fig. 5F) and is not seen thereafter. The MIA-responsive increase of tRF-3s and decrease of 5’ tRNA halves was further confirmed by qRT-PCR (Fig. S9C).

## Discussion

In this study we profiled non-coding small RNAs in three mouse tissues, fetal brain, fetal liver and placenta/decidua and found high levels of tRNA-derived fragments. The tissue specific expression pattern of tRFs was comparable to the differences seen with microRNAs. This lends further credence to the hypothesis that tRF expression or stability is regulated, and that by regulating gene expression and other cellular functions, the tRFs may have a biological role.

We particularly note the high levels of tRNA halves in the mice placenta/decidua as detected by SybrGold staining (Fig. 1B), Northern blots (Fig. 2E) and small RNA-seq (Fig. 1 and Fig. 2). We made relative comparison of same tRFs across different conditions by small RNA-seq, which has been shown to be sensitive to terminal and internal modifications on tRFs. Our small RNA-seq protocol relies on the presence of 5’ phosphate (5’ P) and 3’ hydroxyl (3’ OH) group on the RNA of interests for the adaptor ligation step, therefore any RNAs that harbor terminal modifications such as 3’ cyclic phosphate (3’ cP) /phosphate (3 ‘P) or 5’ hydroxyl (5’ OH) group as direct products of certain ribonucleases [61] or 3’ aminoacylation as shown for 3’ tRNA halves [42] are expected to be under-represented. tRNA halves generation in mammalian cells has been attributed to an RNase A member Angiogenin (ANG), that leaves products with 3’ cP and 5’ OH [61], though recently it has been shown that ANG is not the only enzyme responsible for generating tRNA halves in cell lines [56]. Based on the increase in qRT-PCR signal with T4 PNK treatment (Fig. 3E), we estimate the clonable 5’ tRNA halves detected in small RNA-seq represent ∼ 10-50% of all 5’ tRNA halves in the samples (Fig. 3E). The cloned 5’ tRNA halves could have been generated by other mechanisms that do not generate 3’ cP/3’ P, or could have been converted from a 3’ cP-containing precursor by cellular enzymes in the cell or during extraction. One such cellular enzyme could be the abundant alkaline phosphatase in placenta, that has the ability to convert 3’ P to 3’ OH. Detection of tRFs by RT (reverse transcription)-based approaches could also be affected by internal modifications such as 1-methyladenosine (m1A), which could lead to a combination of read-through (wild-type sequence), mismatch and stopped (truncated) RT products [62]. Only the stopped RT product will not be sequenced in our small RNA-seq protocol (due to lack of 5’ adaptor), which could lead to under-representation of the m1A-containing tRFs. Nevertheless, under the assumption that tRF clonability is not altered by different conditions, one can still make relative comparision by small RNA-seq, which has the advantage of high-throughput, high sensitivity and base resolution over hybridization-based approaches (such as Northern blots and microarray).

Placental/decidual tRNA halves are regulated over time (Fig. 4) and show a transient response to MIA (maternal immune activation) (Fig. 5), suggesting that there may be underappreciated roles of tRNA halves in the interaction between fetus and maternal immune system. tRNA halves have been shown to play diverse roles in different aspects of biology [26, 27]. In particular, tRNA halves can be generated in response to diverse stimuli including viral infection and sex hormone [42, 53, 54, 63] and have been proposed to be new signaling molecules in immune biology [46]. Moreover, tRNA halves were found abundantly in body fluids, including blood/serum, sperm, cerebrospinal fluid and more [63-70]. tRNA halves in mouse sperm were shown to be regulated by parental diet and reported to modulate offspring development [19, 41, 45]. Our finding that high levels of tRNA halves exist in the mouse placenta/decidua is consistent with recent reports about abundant tRNA halves in the human placenta [71, 72]. Why placenta has such high levels of tRNA halves remains poorly understood, but it has been suggested that an oxidative stress environment is generated during gestation [73, 74], and that oxidative stress can trigger tRNA half formation [47, 48, 56]. Here we have focused on the placenta/decidua from fetal development window between E12.5 to E18.5 as this is the window for initiating MIA-related autism phenotypes, and future investigation might reveal whether 5’ tRNA halves are also altered at earlier time points post-implantation. We also noted the sex hormone-dependent tRNA-derived RNAs (SHOT-RNAs), 5’ halves from tRNA^His^ and tRNA^Asp^ [42], are also detected in the maternal-fetal interface (Fig. 2E and Fig. S3C) and are responsive to fetal development or MIA (Fig. 4 and Fig. 5). In particular, 5’ half from tRNA^His^ carries the -1G modification as previously described [75]. It is possible sex hormones might play a role in the biogenesis and function of these placental/decidual tRNA fragments. Intriguingly, tRNA halves were even more enriched in the syncytiotrophoblast-derived extracellular vesicles (STB-EV) that are released from placenta into the maternal circulation [71]. Whether MIA-responsive tRNA halves could travel through maternal circulation or fetal circulation to impact fetal brain maturation and cause neurodevelopmental disease awaits future investigation.

Specifically, 5’ tRNA halves from multiple tRNAs (including but not limited to tRNA^Glu^ and tRNA^Gly^) were down-regulated from E12.5 to E18.5 (Fig. 4) and were down-regulated by MIA (Fig. 5C); while 5’ half from tRNA^His^ was up-regulated over time (Fig. S4) and by poly(I:C)-induced MIA (Fig. S9B). The latter is consistent with the induction of a 5’ half from tRNA^His^ by poly(I:C) treatment on cell lines through RNaseL action [76], an enzyme that is responsive to bacteria and virus infection [77-79] and has been suggested to balance between neuroprotective and neurotoxic functions of microglia [80]. Since tRNA half levels can be regulated by tRNA level, tRNA cleavage, tRNA modification and tRNA half stability [81-86], it will be interesting to examine how each factor contributes to the regulation observed in the maternal-fetal interface.

The 5’ tRNA^Glu^ and tRNA^Gly^ halves in mouse sperm have been shown to regulate transcripts associated with endogenous retroelement MERVL [41], so the placental tRNA halves may repress MERVL elements and thus affect placental function and embryo development. Similarly, 18-nucleotide and 22-nucleotide tRF-3s (altered by MIA, further discussed below) have been shown to inhibit LTR-retrotransposon element activity [44]. Since the previous paper focused on Transposable Element (TE) regulation by tRF-3s from tRNA^Lys^ [44], it is not yet clear whether tRF-3s from other tRNAs, such as the MIA-responsive ones from tRNA^Tyr^, tRNA^Ala^, tRNA^Thr^, tRNA^Gln^, tRNA^Trp^, tRNA^Ser^ and tRNA^Leu^ (Fig. 4C), would also have similar TE regulatory activities. This is particularly interesting given retrotransposon element is important for embryo development, placenta function and evolution [87, 88].

The 18-22 nucleotide long tRF-3s could regulate placenta function or embryo development through protein translation control or/and gene expression regulation. Some tRF-3s can associate with Argonaute proteins and repress mRNA and protein expression by recognizing target mRNAs with complementarity to the tRF-3 sequence [35, 39, 57, 58]. This microRNA-like action could be an important mechanism by which the tRF-3s altered in response to MIA may regulate gene expression in the maternal-fetal interface, or in other fetal or maternal tissues, if transported to the latter.

Several placental/decidual microRNAs are regulated over time during fetal development or in response to MIA, and they could have phenotypic effects by similar mechanisms as tRF-3s. It was thus interesting that the predicted targets of the microRNAs we detected in the placenta are enriched in pathways involved in development. Mmu-miR-215-5p is one of the most up-regulated placental/decidual small RNAs over time from E12.5 to E18.5 (Fig. 4D). miR-215 expression in maternal circulation has been found to correlate with birth-weight-for-gestational-age [89] and has neuroprotective role by regulating IL-17 pathway [90]. Mmu-miR-146b-5p is one of the most down-regulated placental/decidual small RNAs over time (Fig. 4F). miR-146 has been detected in early pregnancy decidua [91] and has important roles in innate immunity [92-94]. We also found miR-291a/b-5p significantly up-regulated by maternal immune activation (Fig. 5C). miR-291a and miR-291b are part of the mESC-specific miR-290-295 cluster, which has been shown to play important roles in early embryo development and maternal-fetal transport [95-97]. Target prediction shows mmu-miR-215-5p, mmu-miR-146b-5p and mmu-miR-291-5p have the potential to regulate development-related pathways (Fig. S10).

The very transient response in the placental/decidual short RNA repertoire after MIA strongly suggests that it might be modulated by maternal cytokines activated by MIA. Poly(I:C) injection induces up-regulation of pro-inflammatory cytokines such as IL-6 in the maternal circulation within several hours after injection [98-100] and similar up-regulation of maternal IL-6 in the placenta [17], which is consistent with the duration of the short RNA response. It is yet to be determined whether the effect of MIA-responsive small RNAs on the transcriptome or proteome will be similarly transient. Alternatively, changes in gene expression initiated by a transient change in a short RNA can continue to be amplified over time even if the initial short RNA changes disappear.

Given the strong male-specific bias in the appearance of autism like phenotypes in the progeny following MIA [11, 12], we were interested in whether there were sex-specific differences in the basal levels or the MIA-induced alterations of tRFs or microRNAs. No tRFs or microRNAs display sex-specific expression at any time point during normal gestation from E12.5 to E18.5 (Fig. S6, FDR < 0.05). The changes in the short RNA repertoire in response to MIA appear not to be affected by the sex of the fetus as determined by likelihood ratio test (described in methods). Since our analysis was done on bulk tissue containing both maternal-derived decidua and fetal-derived placenta, we might have missed small RNA expression difference that is only present in specific cell types. Of course, mRNA repertoires are expected to be different between sexes, and so we cannot rule out a role of the short RNAs on the development of the autism like phenotype. This is because the phenotype ultimately depends on changes in gene expression, and the same short RNAs may have very different effects on gene expression depending on which mRNAs are expressed and susceptible to regulation by the short RNAs.

In summary, by showing tissue-specific, developmentally regulated and MIA responsive changes in the short RNA-repertoire, this study sets the stage for understanding whether and how tRFs, independently or in co-operation with microRNAs, produce the neurodevelopmental changes that lead to autism like behaviors. By discovering the very high levels of tRNA-halves in the placenta, the study also raises questions about whether and how the tRNA-halves might regulate the fetal-maternal interface.

## Materials and methods

### Mice

All mouse experiments were performed in accordance with the relevant guidelines and regulations of the University of Virginia and approved by the University of Virginia Animal Care and Use Committee. C57BL/6 mice were obtained from Taconic Biosciences. Mice were housed in specific pathogen-free conditions.

### Maternal immune activation

Mice were treated similarly as previously described to trigger robust MIA [11, 101]. Mice were mated overnight and the presence of a vaginal plug was designated as E0.5. Each pregnant dam was weighed and administered 20 mg/kg Poly(I:C) potassium salt (Millipore Sigma) or saline by i.p. injections on both E11.5 and E12.5. Embryonic tissues were harvested for individual fetal mouse at indicated time points relative to the E12.5 injection and snap frozen for storage at −80°C. DNA was extracted from embryo tails to determine fetal sex by PCR [102]. RNA was purified from homogenized tissues with Trizol Reagent (Invitrogen) according to manufacturer’s protocol.

### Northern blot analysis

Northern blot was performed as previously [56]. 5 μg Trizol-purified total RNA were resolved on 10% TBE-Urea Novex gel (Thermo Fisher) and then transferred to Amersham Hybond-N+ membrane (GE life sciences). After UV crosslinking and baking, the membrane was blocked by ExpressHyb Hybridization Solution (Takara Bio) and probed with 5’ biotinylated DNA probe at 40°C. Hybridized membrane was detected with Chemiluminescent Nucleic Acid Detection Module Kit (Thermo Fisher). Probe sequences see Table S4.

### T4 PNK treatment and RT-PCR

1 μg total RNA was treated with T4 PNK (NEB) in the presence of 10 mM ATP at 37 °C for 40 minutes and then cleaned up by ethanol precipitation. No treatment control was performed in parallel without adding the enzyme. Cleaned up RNA was then ligated to 3’ pre-adenylated DNA adaptor (NEB, sequence see Table S4) by T4 RNA ligase 2, truncated (NEB) and 5’ RNA adaptor (NEB, sequence see Table S4) by T4 RNA ligase 1 (NEB). cDNA was generated by ProtoScriptII reverse transcriptase (NEB) primed by primer SR-RT (NEB, sequence see Table S4) complementary to the 3’ DNA adaptor. qPCR was performed with QuantiTect SYBR Green PCR Master mix (Qiagen) using primer sets designed to target junctional sequence covering RNA of interest and the adaptor sequence. Amplification efficiency of primer sets was checked by standard curve analysis; specificity of primer sets was checked by melting curve analysis and amplicon size on 8% TBE-PAGE gel. Primer sequences see Table S4. Mmu-miR-26a-5p and mmu-miR-21a-5p were selected as internal controls based on small RNA-seq.

### Small RNA-seq library

Small RNA-seq libraries were prepared similarly as described in [56]. Briefly, RNA quality was first checked by Agilent 4200 TapeStation using High Sensitivity RNA ScreenTape at Genome Analysis and Technology Core (GATC) of University of Virginia, School of Medicine (Fig. S1). Total RNA of RIN (RNA integrity number) more than 7 is subject to NEBNext Small RNA library kit (NEB). After final PCR amplification with indexed-adaptors, each library was run on 8% TBE-PAGE gel to enrich for 15-50 nucleotide insert. A brief workflow and representative gels before size selection are shown in Fig. 1A and Fig. 1B. Individual libraries were purified from gel and pooled for sequencing on Illumina NextSeq 500 sequencer with high-output, 75-cycles single-end mode at University of Virginia GATC core. The data is deposited to GEO with accession GSE139191 (including 84 samples as detailed in Table S1).

### Data analysis

Small RNA-seq analysis was performed similarly as described in [56] with slight modifications. Cutadapt v1.15 [103] was used to trim 3’ adaptor sequence and remove read length shorter than 15 nt. The reads were then mapped to mouse genome (gencode release M23, GRCm38.p6 ALL) with corresponding annotation plus predicted tRNA genes by STAR aligner v2.5.4 [104]. In general, each library has 5-20 million mapped reads. Total number of mapped reads and percentage of mapping is summarized in Table S1. To quantify small RNAs, unitas v1.7.3 [105] (with SeqMap v1.0.13 [106]) was used to map the reads to mouse sequence of miRBase Release 22 [107], genomic tRNA database [108], Ensembl Release 96 and SILVA rRNA database Release 132. Unitas setting (– species_miR_only –species mus_musculus) was used. In general, we allowed 2 non-templated 3’ nucleotides and 1 internal mismatch for miRNA mapping, and 1 mismatch and 0 insertion/deletion for tRNA fragments mapping (equivalent to – tail 2 –intmod 1 –mismatch 1 –insdel 0). microRNA reads were grouped by mature microRNA names, tRF reads were grouped by parental tRNAs and tRF types (cutoff for tRNA halves is 30 nucleotide long); in the case of multi-mapping, a read was counted as fraction to avoid duplicate counts. For sequence-level analysis (for Figure 3 and Figure 4), we used more stringent mapping without allowing for mismatch (equivalent to –quick).

For differential analysis, DESeq2 (v1.24.0) [109] was used on count matrix of tRFs and miRs. To normalize the data by library size, size factor for each library was estimated using median ratio method [110]. To better estimate log2foldchange for visualization in MA plots and volcano plots, the log2foldchange value from DESeq2 was shrunk using apeglm method [111]. Principle component analysis was done on DESeq2 rlog (regularized log) transformed count matrix on unique sequence level and visualized as two-dimensional plots. For sequence-level analysis (Fig. 3 and Fig. 4), count matrix was built for each unique sequence, with at least 10 read counts in all samples; otherwise, no pre-filtering was done. To identify dynamically expressed miRs and tRFs, control (saline treated) samples at 3 hour, 24 hour, 48 hour and 144 hour time points were used for E12.5, E13.5, E14.5 and E18.5 correspondingly. Heatmap in Fig. 3B was generated using mean reads per million mapped reads value per time point with heatplot function (default setting) in made4_1.58.0 [112], with expression cut-off of 100 RPM and time-dependent change was selected for presentation. To test for any potential effect from fetal sex, we first considered all three factors (1. fetal sex, 2. time point and 3. treatment) and all possible interactions among the three factors (1+2, 1+3, 2+3 and 1+2+3) during differential analysis using DESeq2 on the whole dataset (12 hour time point is omitted due to small sample size). We then performed likelihood ratio test after removing certain factors to see whether considering those factors (or interactions) improve the differential analysis. By this method, it is clear that considering the triple interaction term, or the interaction term between fetal sex and treatment or the interaction term between fetal sex and time point do not improve the model (no p.adj < 0.1); this suggests there are no small RNAs that displayed sex-specific responses to MIA at any time point, and there is no small RNAs that displayed sex-specific expression during development (consistent with Fig. S5). These R/Bioconductor packages (R version 3.6.1) were used for data visualization: ggplot2_3.2.1, ggthemes_4.2.0, dplyr_0.8.3, readr_1.3.1, ggalt_0.4.0, ggpubr_0.2.3, RColorBrewer_1.1-2.

For microRNA target prediction, TargetScanMouse (release v7.2) [113] was used to obtain predicted microRNA target genes based on seed matches and conservation. Predicted genes were submitted to PANTHER [114] overrepresentation test (released 20190711) for GO Ontology database [115] (released 20190703) for mouse biological process against all mouse genes as reference list with default setting (Fisher’s exact Test with Benjamini-Hochberg False Discovery Rate correction).

## Acknowledgement

We would like to thank University of Virginia GATC (genome analysis and technology core) for NGS service and Research Computing Core for computational support.

