## Supplementary material for "tRNA-derived fragments and microRNAs in the maternal-fetal interface of a mouse maternal-immune-activation autism model": Fig. S, Table S

#### *Supplementary figures*

**Figure S1.** Representative TapeStation RNA QC results for RNA from three tissue types.

**Figure S2.** Small RNA profiling in three tissue types (E13.5).

**Figure S3.** 5' and 3' tRNA fragments in three tissue types (E13.5).

**Figure S4.** Detection of 3' tRNA fragments in placenta/decidua.

**Figure S5**. Composition of tRF types and parental tRNAs in E13.5 tissues.

**Figure S6.** Lack of differential expression in placental/decidual small RNAs by fetal sex.

**Figure S7.** Dynamic expression of placental/decidual tRNA fragemnts.

**Figure S8.** Acute response in placental/decidual small RNAs by MIA.

**Figure S9.** MIA-responsive small RNA expression changes at the maternal-fetal interface.

**Figure S10.** Over-represented development-related pathways for the predicted targets of dynamic or MIA-responsive placental microRNAs.

#### *Supplementary tables*

**Table S1.** Mapping statistics of small RNA-seq.

**Table S2.** Dynamically expressed placental/decidual microRNA and tRF sequences.

**Table S3.** MIA-responsive placental/decidual microRNA and tRF sequences from DESeq2.

**Table S4.** Probe/primer sequences.

**Fig. S1**

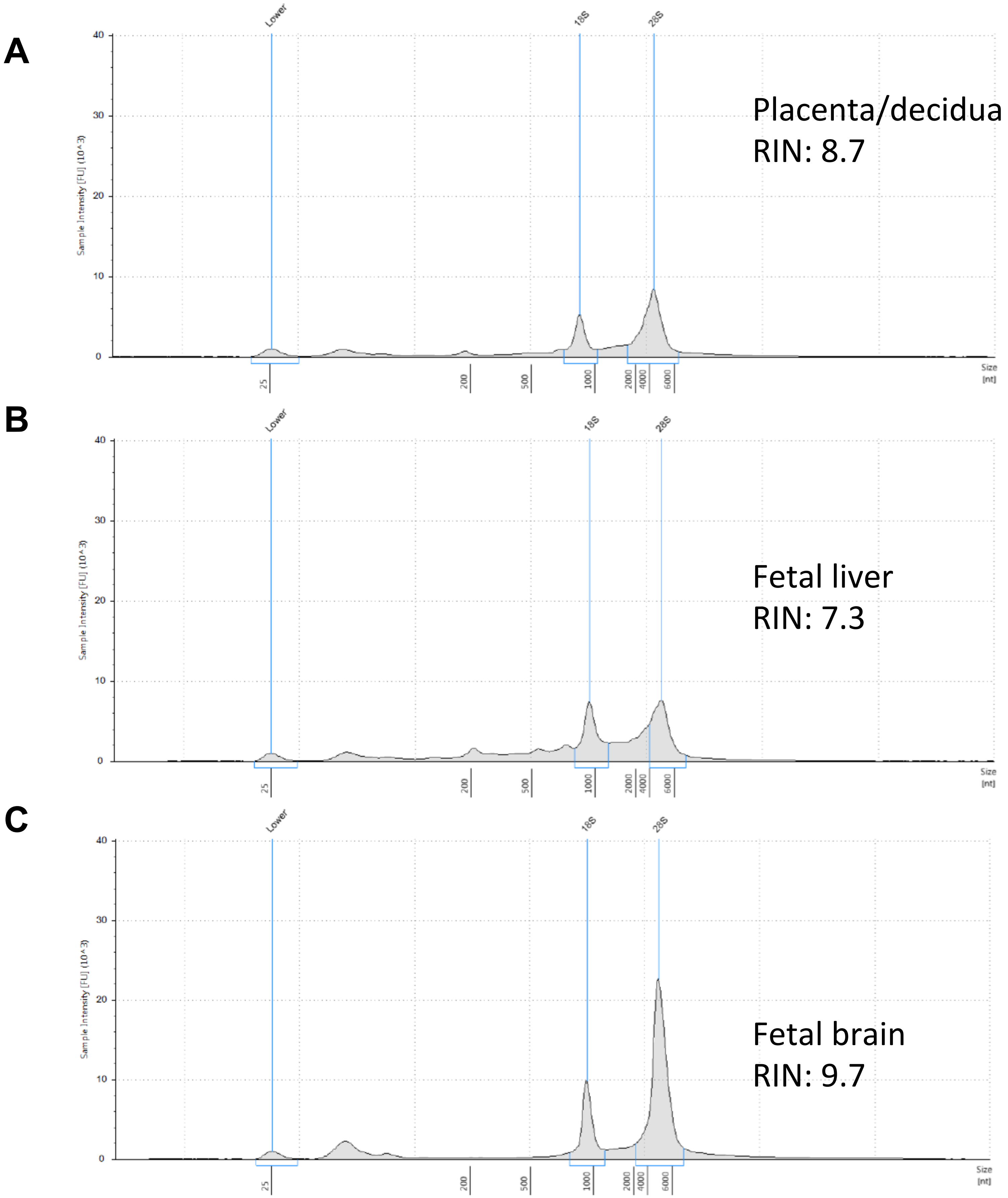


**Figure S1. Representative TapeStation RNA QC results for RNA from three tissue types.** Y-axis: sample intensity (FU-fluorescence unit), X-axis: size (nucleotide). RIN (RNA integrity number) more than 7 was used for small RNA NGS library construction.

Fig. S2

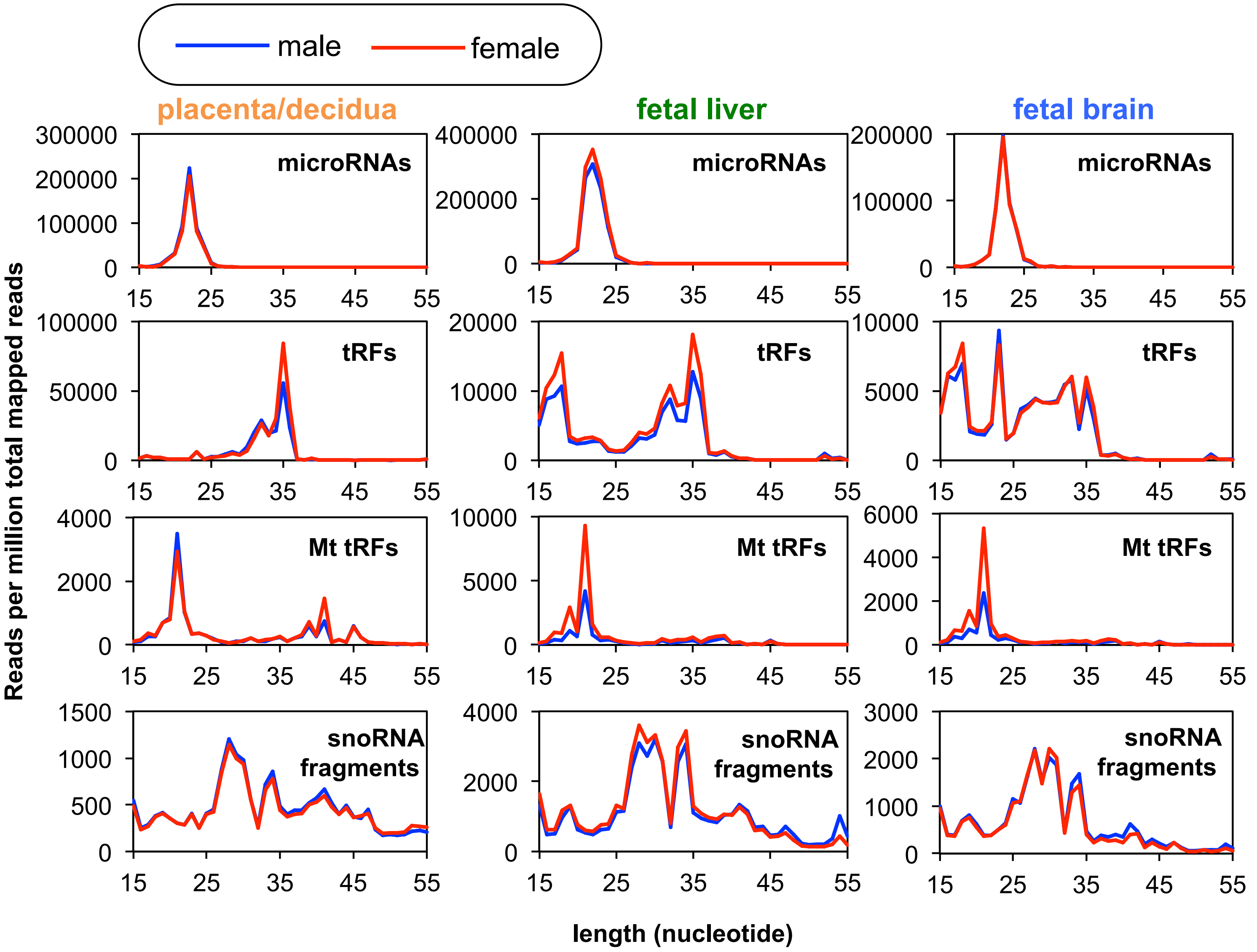


**Figure S2. Small RNA profiling in three tissue types (E13.5).**
Length distribution of microRNAs, tRFs and mitochondria tRFs in three tissue types (E13.5 placenta/decidua, fetal liver and fetal brain) divided by fetal sex (blue: male, red: female, n = 3). X-axis: fragment length (nucleotide), Y-axis: average reads per million total mapped reads.

**Fig. S3**

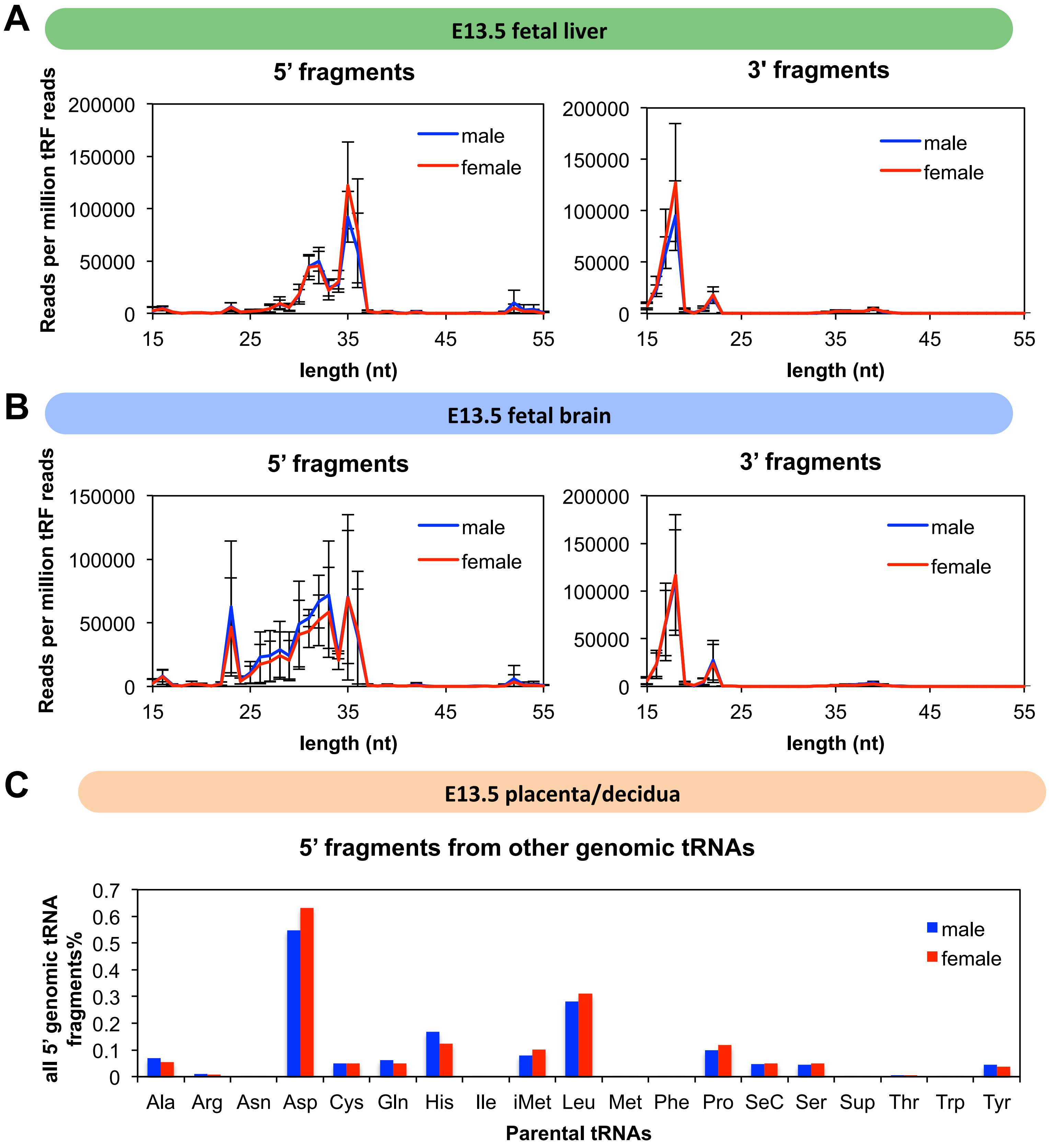


**Figure S3. 5' and 3' tRNA fragments in three tissue types (E13.5).**
(A-B) Length distribution of reads that map to 5' end (left) and 3' end (right) of tRNA (Y-axis: average reads per million total tRF reads, X-axis: fragment length in nucleotide) in E13.5 fetal liver (A) and fetal brain (B). Samples are divided by fetal sex (blue: male, rad: female, n = 3). (C) 5' tRNA fragments from indicated parental tRNAs (excluding the four abundant $tRNA^{Gly}$, $tRNA^{Glu}$, $tRNA^{Val}$ and $tRNA^{Lys}$), corresponding to Fig. 2C. Y-axis: relative percentage among all 5' tRFs.

### Fig. S4

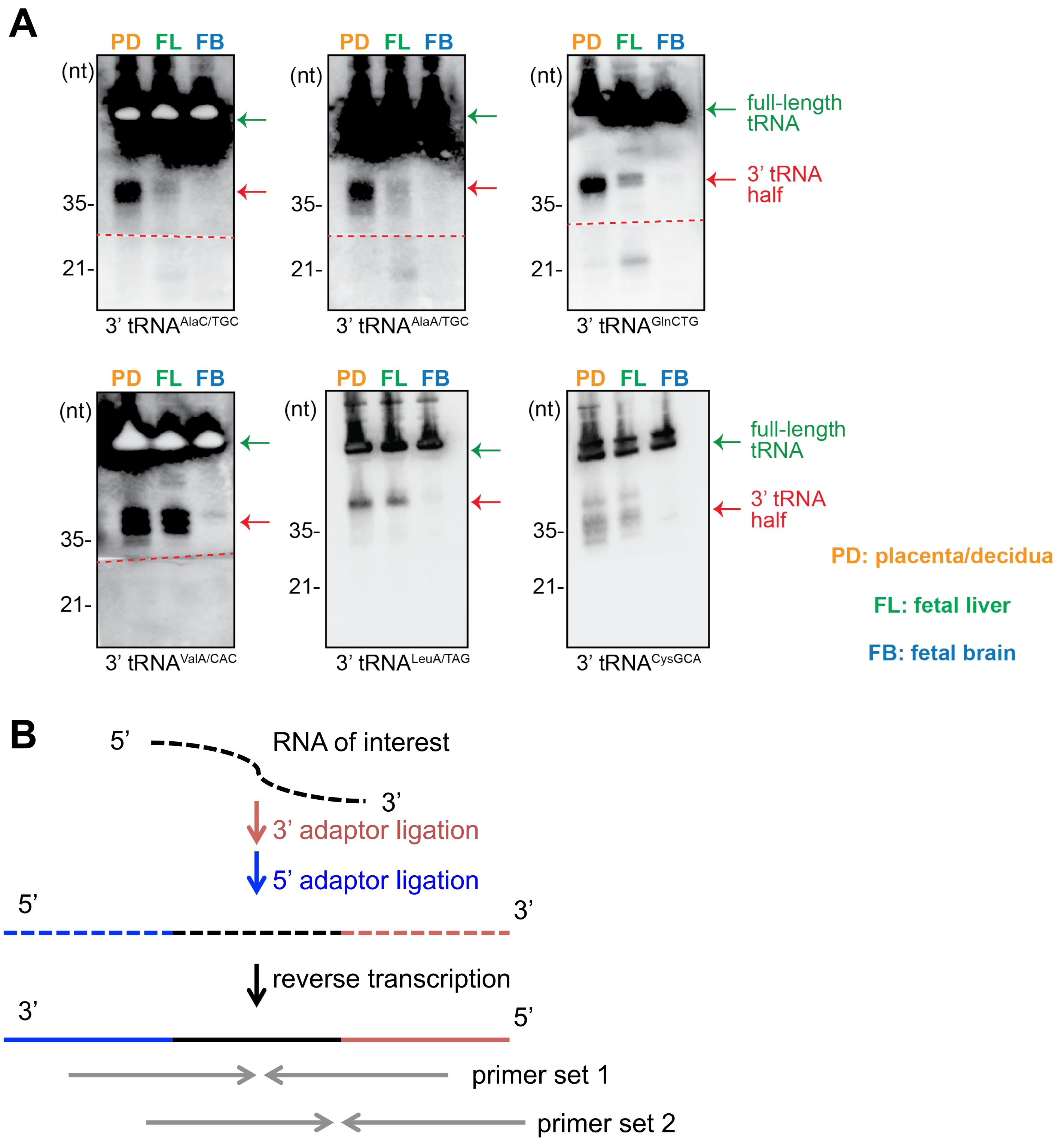


**Figure S4. Detection of 3’ tRNA fragments in placenta/decidua.**
(A) Northern blot detection of 3’ tRNA fragments in normal E13.5 tissues (PD: placenta/decidua, FL: fetal liver, FB: fetal brain). Arrows indicate full-length tRNAs (green arrows) and 3’ tRNA halves (red arrows).
(B) Scheme of the modified qRT-PCR method for tRF detection. RNA of interest is ligated to 3’ adaptor and 5’ adaptor and subjected to reverse transcription. Primers target the junctional sequence will specifically detect tRFs.

#### Fig. S5

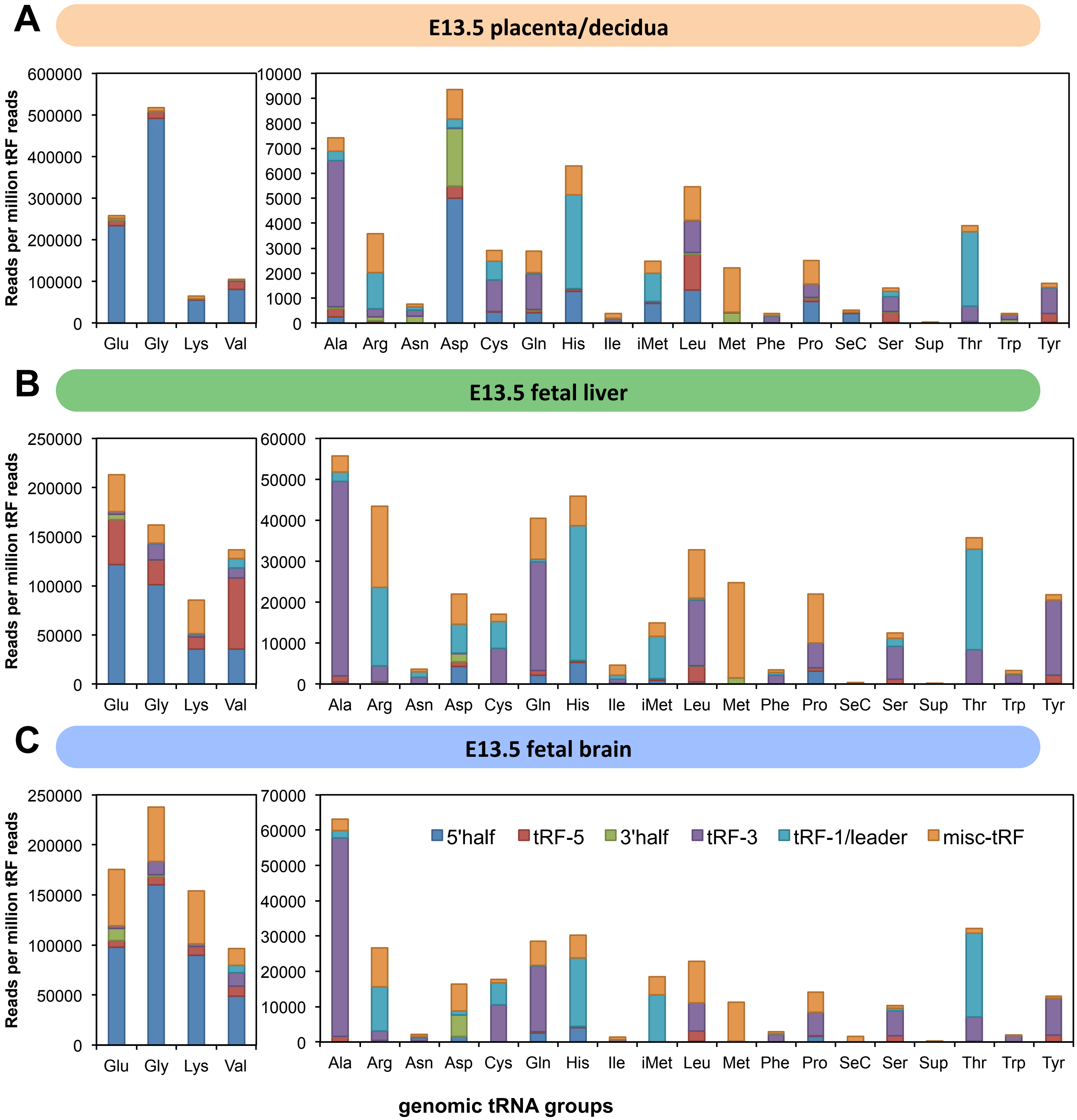


**Figure S5. Composition of tRF types and parental tRNAs in E13.5 tissues.**
(A-C) Detailed composition of tRF types and parental tRNAs in E13.5 placenta/decidua (A), fetal liver (B) and fetal brain (C). Samples from both fetal sex are combined (n = 6). Y-axis: reads per million total tRF reads (averaged from 6 samples). X-axis: parental tRNAs where tRFs are mapped. Definition of tRF types see main text and methods section.

**Fig. S6**

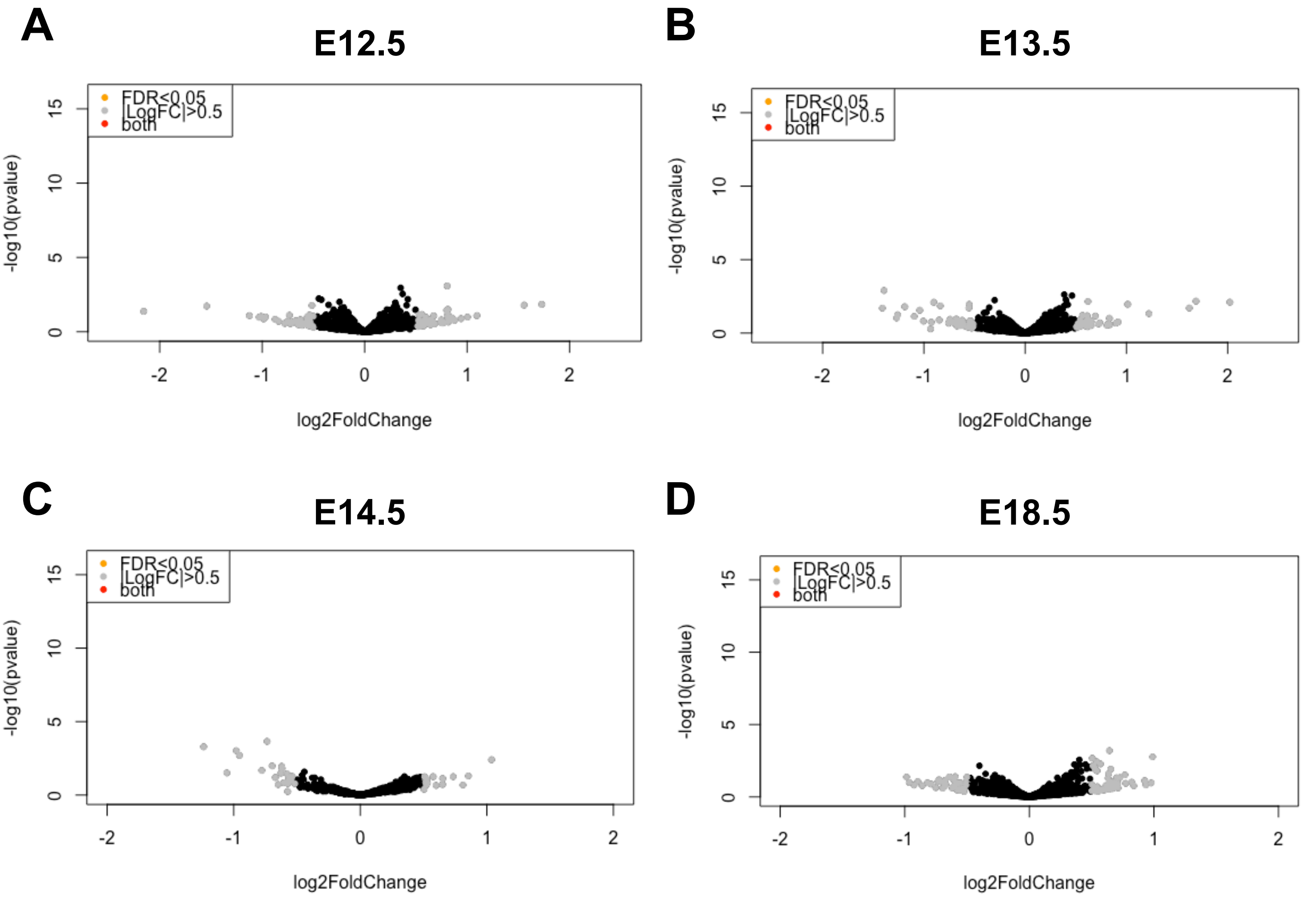


**Figure S6. Lack of differential expression in placental/decidual small RNAs by fetal sex.** (A-D) Differential analysis was performed on the union of microRNAs and tRFs for placenta/decidua samples from each time point (E12.5, E13.5, E14.5 and E18.5) separately between fetal sex. Volcano plots: X-axis: Log2FoldChange (female versus male); Y-axis: -log10 (p value); FDR (false discovery rate) of 0.05. Significant changes (Log2FoldChange more than 0.5 or less than -0.5, and p adjusted less than 0.05) will show as red dots. E12.5/13.5/14.5: n = 6 (3 males and 3 females); E18.5: n = 8 (4 males and 4 females).

**Fig. S7**

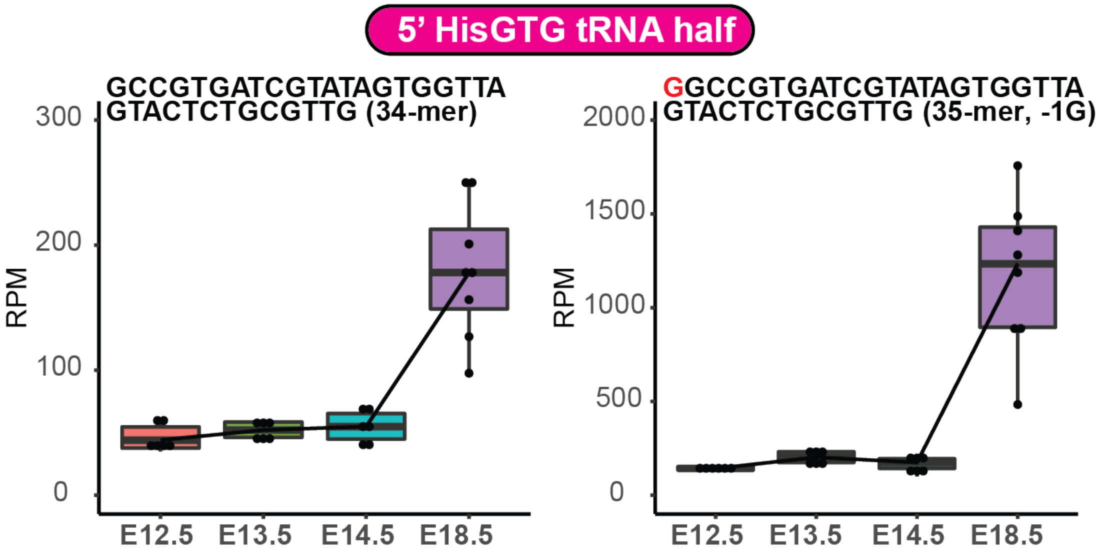


**Figure S7. Dynamic expression of placental/decidual tRNA fragments.**
5' half from tRNA-His-GTG shows temporal increase in placenta/decidua from E12.5 to E18.5 in small RNA-seq (RPM: reads per million mapped reads).

### Fig. S8

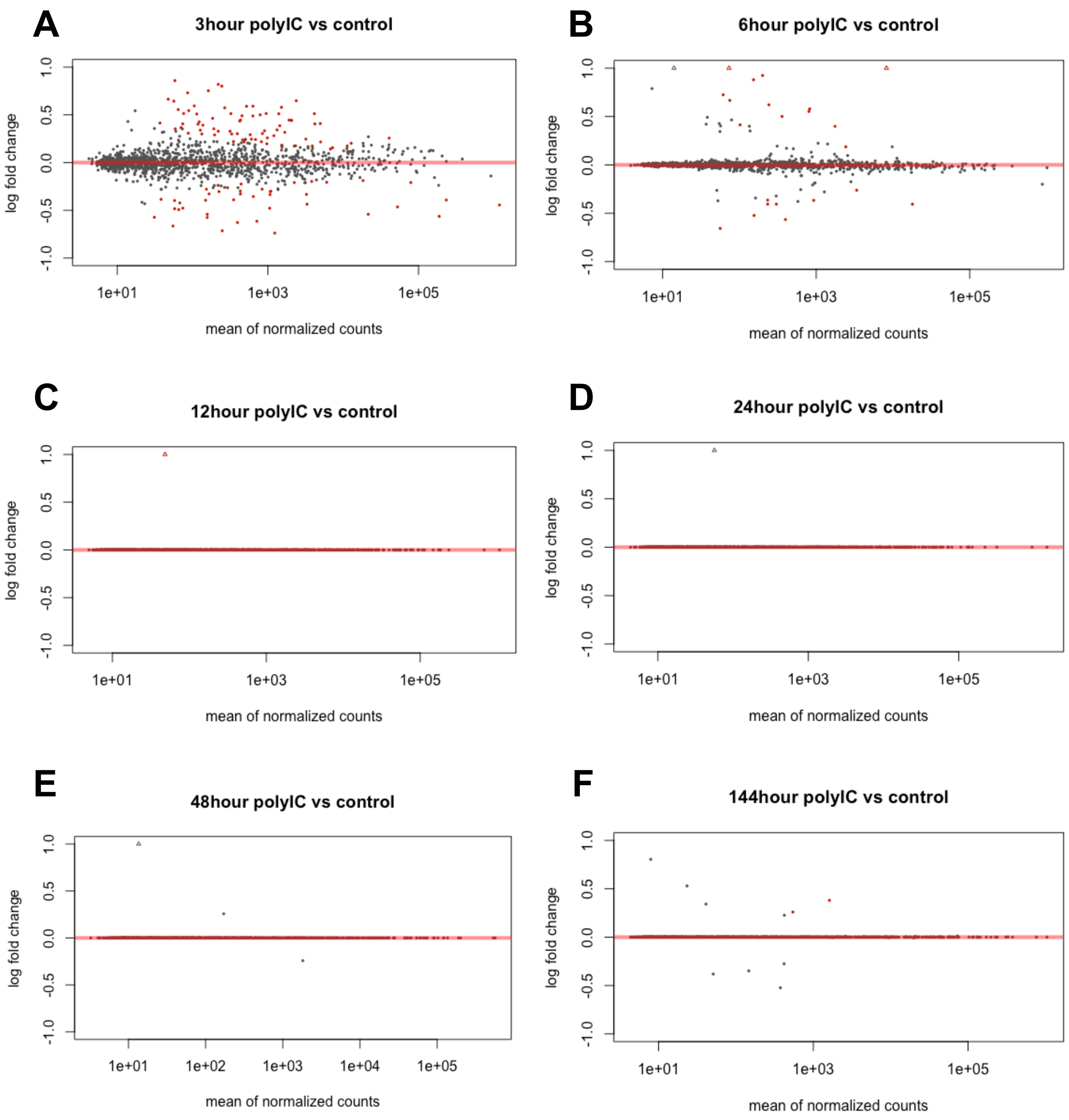


**Figure S8. Acute response in placental/decidual small RNAs by MIA.**
(A-F) Differential analysis was performed on the union of microRNAs and tRFs for placenta/ decidua samples from each time point between MIA and control (fetal sex combined, n = 4 for 12 hrs, n = 8 for 144 hrs and n = 6 for all the other time points). MA plots: X-axis: mean of normalized counts; Y-axis: log2 fold change (MIA/control). Significant changes (p adjusted less than 0.05) will show as red dots; triangles denote changes beyond Y-axis limit (more than 2-fold).

Fig. S9

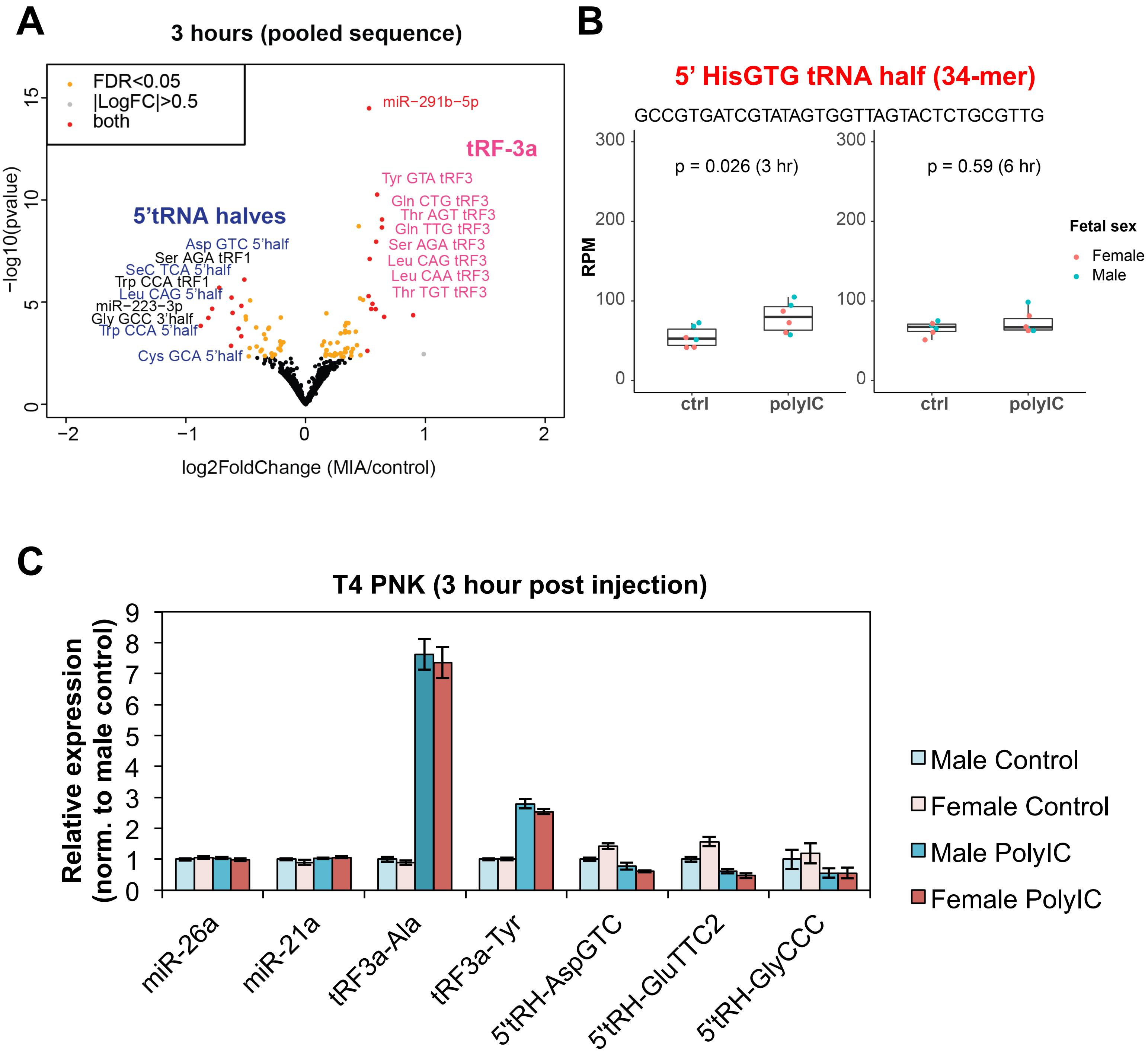


**Figure S9. MIA-responsive small RNA expression changes at the maternal-fetal interface.** (A) Volcano plot of MIA-dependent changes at 3 hour time point (similar as Fig. 4C, on pooled sequence level). Significant miRs and tRFs (adjusted $p < 0.05$) are labeled. (B) Box plots showing RPM (reads per million total mapped reads) for specific miR or tRF sequence, with each dot represents one sample (n = 6 for each time point, fetal sex is specified by dot color) and middle line represents median value. To compare the difference between control group and poly(I:C) (MIA) group, unpaired two-samples Wilcoxon test was performed (n = 6 for each group). (C) qRT-PCR detection of MIA-responsive changes in tRF-3s and 5' tRNA halves. Error bars represent standard deviation from 3 biological replicates at 3 hour time point. Relative expression was calculated by dCt normalized to male control sample.

#### Fig. S10

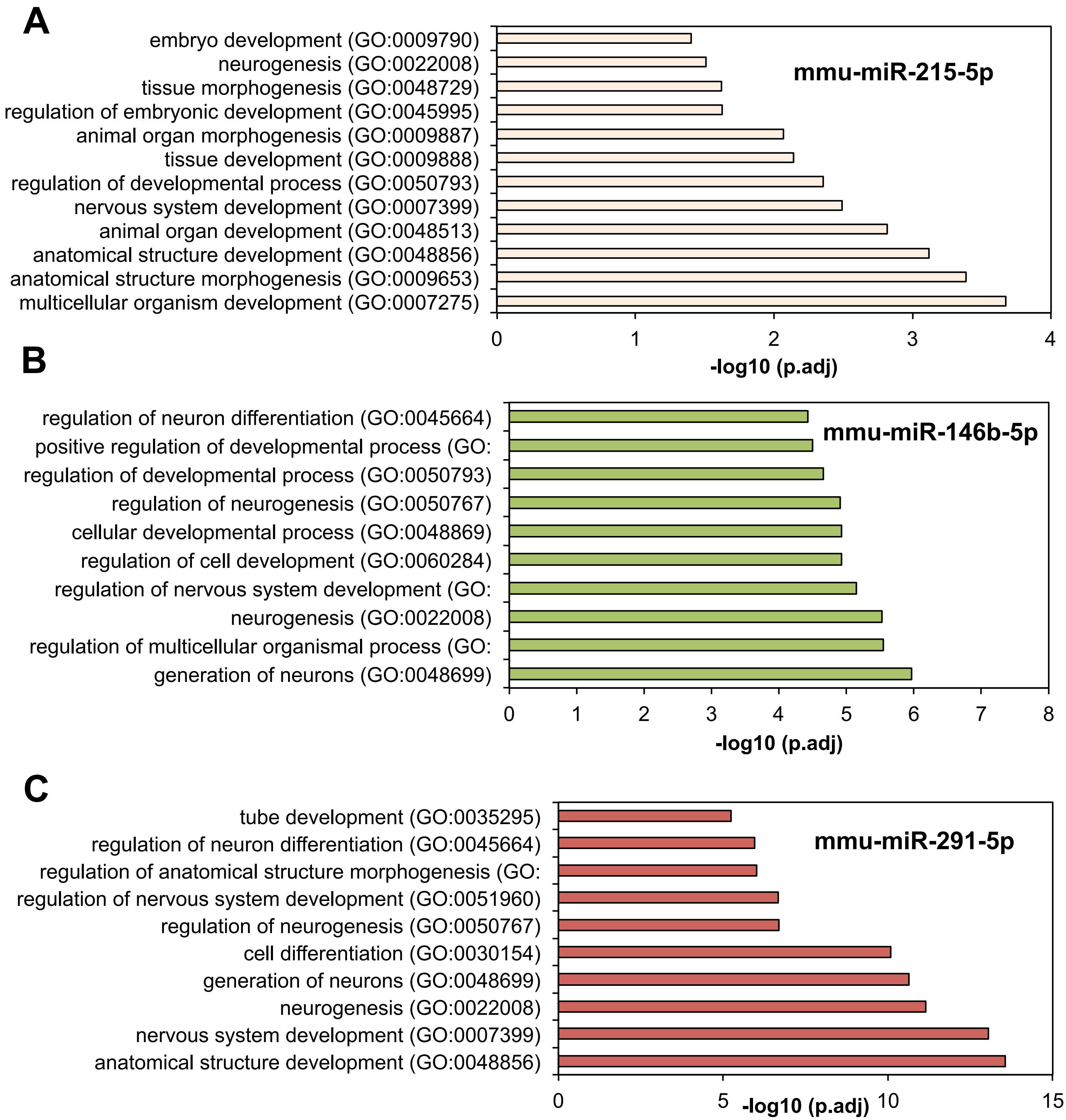


**Figure S10. Over-represented development-related pathways for the predicted targets of dynamic or MIA-responsive placental microRNAs.**

(A-C) Over-represented development-related biological pathways of the TargetScan predicted targets of specific placental microRNAs identified from Fig. 3 (dynamically expressed) and Fig. 4 (MIA-responsive).

**Table S1. Mapping statistics of small RNA-seq (GSE139191)**

| GEO number | Used in Figure # | Condition | Total Reads | Reads with adaptors | After cutadapt (>=15nt) | Average insert length | Mapping % (STAR aligner) |
|---|---|---|---|---|---|---|---|
| GSM4133305 | 1, 2 | E13.5 fetal brain male control1 | 12,004,154 | 11,861,370 | 8,405,397 | 23 | 94.56 |
| GSM4133306 | 1, 2 | E13.5 fetal brain female control1 | 13,695,073 | 13,496,385 | 9,093,162 | 24 | 94.59 |
| GSM4133307 | 1, 2 | E13.5 fetal brain male control2 | 12,074,541 | 11,973,383 | 9,503,235 | 26 | 94.26 |
| GSM4133308 | 1, 2 | E13.5 fetal brain female control2 | 11,137,835 | 11,012,387 | 7,415,973 | 24 | 92.92 |
| GSM4133309 | 1, 2 | E13.5 fetal brain male control3 | 194,922,456 | 193,330,397 | 172,760,436 | 27 | 98.85 |
| GSM4133310 | 1, 2 | E13.5 fetal brain female control3 | 77,981,100 | 77,214,001 | 67,326,209 | 26 | 98.73 |
| GSM4133311 | 1, 2 | E13.5 fetal liver male control1 | 11,117,222 | 11,020,519 | 10,371,805 | 28 | 97.94 |
| GSM4133312 | 1, 2 | E13.5 fetal liver female control1 | 10,985,020 | 10,886,377 | 10,148,049 | 28 | 97.83 |
| GSM4133313 | 1, 2 | E13.5 fetal liver male control2 | 21,134,992 | 21,024,941 | 18,362,864 | 23 | 97.15 |
| GSM4133314 | 1, 2 | E13.5 fetal liver female control2 | 21,034,510 | 20,905,107 | 17,928,170 | 23 | 97.43 |
| GSM4133315 | 1, 2 | E13.5 fetal liver male control3 | 20,724,371 | 20,597,009 | 17,703,497 | 32 | 98.20 |
| GSM4133316 | 1, 2 | E13.5 fetal liver female control3 | 16,074,486 | 15,969,808 | 12,667,992 | 28 | 97.60 |
| GSM4133317 | 3, 4 | E12.5 placenta/decidua male 3hr control1 | 22,155,870 | 21,411,591 | 19,442,642 | 31 | 97.61 |
| GSM4133321 | 3, 4 | E12.5 placenta/decidua male 3hr control2 | 17,314,279 | 17,000,981 | 15,798,654 | 31 | 97.89 |
| GSM4133325 | 3, 4 | E12.5 placenta/decidua male 3hr control3 | 16,776,452 | 16,399,281 | 14,979,961 | 31 | 97.64 |
| GSM4133318 | 3, 4 | E12.5 placenta/decidua female 3hr control1 | 10,570,579 | 10,338,277 | 9,282,870 | 30 | 97.84 |
| GSM4133322 | 3, 4 | E12.5 placenta/decidua female 3hr control2 | 20,713,040 | 20,191,214 | 18,952,799 | 31 | 98.07 |
| GSM4133326 | 3, 4 | E12.5 placenta/decidua female 3hr control3 | 18,193,953 | 17,829,855 | 16,740,063 | 31 | 97.64 |
| GSM4133349 | 1, 2, 3, 4 | E13.5 placenta/decidua male 24hr control1 | 7,069,811 | 6,924,869 | 6,571,164 | 32 | 98.28 |
| GSM4133353 | 1, 2, 3, 4 | E13.5 placenta/decidua male 24hr control2 | 11,305,192 | 10,853,295 | 10,719,465 | 33 | 98.41 |
| GSM4133357 | 1, 2, 3, 4 | E13.5 placenta/decidua male 24hr control3 | 9,592,914 | 9,414,157 | 9,297,569 | 32 | 98.67 |
| GSM4133350 | 1, 2, 3, 4 | E13.5 placenta/decidua female 24hr control1 | 9,586,706 | 9,455,962 | 9,071,012 | 31 | 98.39 |
| GSM4133354 | 1, 2, 3, 4 | E13.5 placenta/decidua female 24hr control2 | 13,163,095 | 12,777,487 | 12,625,757 | 32 | 98.66 |
| GSM4133358 | 1, 2, 3, 4 | E13.5 placenta/decidua female 24hr control3 | 12,264,488 | 11,842,554 | 11,739,325 | 32 | 98.59 |
| GSM4133361 | 3, 4 | E14.5 placenta/decidua male 48hr control1 | 8,630,163 | 8,454,243 | 8,055,667 | 31 | 98.33 |
| GSM4133365 | 3, 4 | E14.5 placenta/decidua male 48hr control2 | 10,821,742 | 10,674,665 | 10,205,397 | 30 | 98.00 |
| GSM4133369 | 3, 4 | E14.5 placenta/decidua male 48hr control3 | 10,571,553 | 10,348,922 | 9,780,795 | 32 | 98.14 |
| GSM4133362 | 3, 4 | E14.5 placenta/decidua female 48hr control1 | 7,852,119 | 7,752,175 | 7,160,119 | 31 | 98.21 |
| GSM4133366 | 3, 4 | E14.5 placenta/decidua female 48hr control2 | 9,905,540 | 9,717,263 | 9,617,499 | 31 | 97.94 |
| GSM4133370 | 3, 4 | E14.5 placenta/decidua female 48hr control3 | 9,138,841 | 8,828,260 | 8,511,796 | 33 | 97.99 |
| GSM4133373 | 3, 4 | E18.5 placenta/decidua male 144hr control1 | 20,724,182 | 20,154,569 | 19,751,586 | 30 | 98.30 |
| GSM4133377 | 3, 4 | E18.5 placenta/decidua male 144hr control2 | 11,320,582 | 11,195,999 | 10,755,727 | 30 | 98.46 |
| GSM4133381 | 3, 4 | E18.5 placenta/decidua male 144hr control3 | 11,821,617 | 11,582,833 | 11,120,279 | 31 | 98.25 |
| GSM4133385 | 3, 4 | E18.5 placenta/decidua male 144hr control4 | 12,234,408 | 11,816,072 | 11,645,757 | 31 | 98.36 |
| GSM4133374 | 3, 4 | E18.5 placenta/decidua female 144hr control1 | 10,306,850 | 10,177,951 | 9,851,943 | 30 | 98.29 |
| GSM4133378 | 3, 4 | E18.5 placenta/decidua female 144hr control2 | 13,599,913 | 13,390,764 | 12,874,738 | 30 | 98.14 |
| GSM4133382 | 3, 4 | E18.5 placenta/decidua female 144hr control3 | 17,560,682 | 17,308,485 | 16,468,200 | 30 | 98.20 |
| GSM4133386 | 3, 4 | E18.5 placenta/decidua female 144hr control4 | 16,700,936 | 16,258,054 | 15,579,790 | 31 | 98.22 |
| GSM4133319 | 4 | E12.5 placenta/decidua male 3hr polyIC1 | 7,252,680 | 7,076,722 | 6,033,179 | 30 | 97.46 |
| GSM4133323 | 4 | E12.5 placenta/decidua male 3hr polyIC2 | 15,142,027 | 14,774,040 | 14,030,147 | 31 | 97.61 |
| GSM4133327 | 4 | E12.5 placenta/decidua male 3hr polyIC3 | 15,036,791 | 14,712,283 | 13,994,867 | 31 | 97.46 |
| GSM4133320 | 4 | E12.5 placenta/decidua female 3hr polyIC1 | 13,155,138 | 12,890,129 | 11,584,431 | 29 | 97.00 |
| GSM4133324 | 4 | E12.5 placenta/decidua female 3hr polyIC2 | 18,532,834 | 17,973,691 | 16,997,963 | 32 | 97.90 |
| GSM4133328 | 4 | E12.5 placenta/decidua female 3hr polyIC3 | 16,193,035 | 15,871,827 | 15,174,315 | 33 | 97.79 |
| GSM4133351 | 4 | E13.5 placenta/decidua male 24hr polyIC1 | 11,975,778 | 11,758,997 | 11,276,894 | 32 | 98.36 |
| GSM4133355 | 4 | E13.5 placenta/decidua male 24hr polyIC2 | 12,385,081 | 11,962,278 | 11,314,378 | 32 | 98.53 |
| GSM4133359 | 4 | E13.5 placenta/decidua male 24hr polyIC3 | 12,740,257 | 12,525,033 | 12,091,557 | 32 | 98.20 |
| GSM4133352 | 4 | E13.5 placenta/decidua female 24hr polyIC1 | 13,404,007 | 13,107,129 | 12,787,715 | 32 | 98.60 |
| GSM4133356 | 4 | E13.5 placenta/decidua female 24hr polyIC2 | 7,713,915 | 7,528,422 | 7,311,759 | 31 | 98.61 |
| GSM4133360 | 4 | E13.5 placenta/decidua female 24hr polyIC3 | 8,701,808 | 8,531,538 | 8,173,924 | 32 | 98.62 |
| GSM4133363 | 4 | E14.5 placenta/decidua male 48hr polyIC1 | 5,827,335 | 5,710,680 | 5,539,973 | 32 | 98.20 |
| GSM4133367 | 4 | E14.5 placenta/decidua male 48hr polyIC2 | 10,017,750 | 9,367,623 | 9,581,262 | 34 | 97.03 |
| GSM4133371 | 4 | E14.5 placenta/decidua male 48hr polyIC3 | 8,672,120 | 8,546,381 | 7,996,963 | 32 | 96.61 |
| GSM4133364 | 4 | E14.5 placenta/decidua female 48hr polyIC1 | 5,850,609 | 5,720,147 | 5,604,939 | 32 | 98.18 |
| GSM4133368 | 4 | E14.5 placenta/decidua female 48hr polyIC2 | 11,244,401 | 10,793,419 | 10,654,123 | 32 | 97.30 |
| GSM4133372 | 4 | E14.5 placenta/decidua female 48hr polyIC3 | 11,841,761 | 11,629,020 | 11,145,412 | 30 | 97.04 |
| GSM4133375 | 4 | E18.5 placenta/decidua male 144hr polyIC1 | 10,886,369 | 10,680,599 | 10,242,050 | 32 | 98.34 |

| | | | | | | | |
|---|---|---|---|---|---|---|---|
| GSM4133379 | 4 | E18.5 placenta/decidua male 144hr polyIC2 | 15,215,327 | 14,888,649 | 14,338,091 | 30 | 98.07 |
| GSM4133383 | 4 | E18.5 placenta/decidua male 144hr polyIC3 | 14,728,000 | 14,483,671 | 14,021,234 | 30 | 98.10 |
| GSM4133387 | 4 | E18.5 placenta/decidua male 144hr polyIC4 | 13,651,934 | 13,269,042 | 13,068,473 | 30 | 98.55 |
| GSM4133376 | 4 | E18.5 placenta/decidua female 144hr polyIC1 | 9,249,512 | 9,092,129 | 8,685,705 | 32 | 98.40 |
| GSM4133380 | 4 | E18.5 placenta/decidua female 144hr polyIC2 | 14,137,555 | 13,817,768 | 13,534,395 | 30 | 98.17 |
| GSM4133384 | 4 | E18.5 placenta/decidua female 144hr polyIC3 | 11,883,745 | 11,694,465 | 11,477,855 | 31 | 98.70 |
| GSM4133388 | 4 | E18.5 placenta/decidua female 144hr polyIC4 | 15,733,121 | 15,441,084 | 14,964,521 | 30 | 98.60 |
| GSM4133329 | 4 | E12.5 placenta/decidua male 6hr control1 | 21,290,545 | 20,432,270 | 18,462,355 | 31 | 97.07 |
| GSM4133333 | 4 | E12.5 placenta/decidua male 6hr control2 | 15,280,249 | 14,971,456 | 14,193,565 | 32 | 97.62 |
| GSM4133337 | 4 | E12.5 placenta/decidua male 6hr control3 | 18,078,322 | 17,685,002 | 15,736,064 | 30 | 96.65 |
| GSM4133330 | 4 | E12.5 placenta/decidua female 6hr control1 | 17,779,714 | 17,232,908 | 15,907,111 | 32 | 97.40 |
| GSM4133334 | 4 | E12.5 placenta/decidua female 6hr control2 | 11,221,487 | 10,839,366 | 10,647,097 | 32 | 97.86 |
| GSM4133338 | 4 | E12.5 placenta/decidua female 6hr control3 | 12,739,960 | 12,466,129 | 11,595,273 | 30 | 97.48 |
| GSM4133331 | 4 | E12.5 placenta/decidua male 6hr polyIC1 | 18,123,829 | 17,532,193 | 16,461,766 | 31 | 97.21 |
| GSM4133335 | 4 | E12.5 placenta/decidua male 6hr polyIC2 | 11,571,942 | 11,356,781 | 10,062,929 | 30 | 97.04 |
| GSM4133339 | 4 | E12.5 placenta/decidua male 6hr polyIC3 | 11,688,220 | 11,373,296 | 10,668,140 | 31 | 97.62 |
| GSM4133332 | 4 | E12.5 placenta/decidua female 6hr polyIC1 | 14,839,382 | 14,468,182 | 13,826,516 | 31 | 97.64 |
| GSM4133336 | 4 | E12.5 placenta/decidua female 6hr polyIC2 | 13,907,684 | 13,619,130 | 11,809,454 | 30 | 96.91 |
| GSM4133340 | 4 | E12.5 placenta/decidua female 6hr polyIC3 | 12,247,573 | 11,929,290 | 10,970,089 | 31 | 97.30 |
| GSM4133341 | 4 | E12.5 placenta/decidua male 12hr control1 | 10,632,095 | 10,491,484 | 9,961,267 | 31 | 98.08 |
| GSM4133345 | 4 | E12.5 placenta/decidua male 12hr control2 | 8,441,882 | 8,286,251 | 7,650,795 | 31 | 98.23 |
| GSM4133342 | 4 | E12.5 placenta/decidua female 12hr control1 | 9,910,445 | 9,675,257 | 9,177,876 | 31 | 98.21 |
| GSM4133346 | 4 | E12.5 placenta/decidua female 12hr control2 | 10,446,918 | 10,181,739 | 9,609,198 | 30 | 98.18 |
| GSM4133343 | 4 | E12.5 placenta/decidua male 12hr polyIC1 | 20,623,773 | 20,249,555 | 19,554,059 | 32 | 97.90 |
| GSM4133347 | 4 | E12.5 placenta/decidua male 12hr polyIC2 | 10,166,684 | 9,919,779 | 9,136,598 | 30 | 98.16 |
| GSM4133344 | 4 | E12.5 placenta/decidua female 12hr polyIC1 | 12,274,296 | 12,117,264 | 11,371,668 | 31 | 98.02 |
| GSM4133348 | 4 | E12.5 placenta/decidua female 12hr polyIC2 | 9,700,714 | 9,564,721 | 8,899,653 | 31 | 98.19 |

**Table S2. Dynamically expressed placental/decidual microRNA and tRF sequences.**

| Type | Length | Sequence | Annotation | E12.5_mean _log10RPM | E13.5_mean _log10RPM | E14.5_mean _log10RPM | E18.5_mean _log10RPM | Decrease/ increase over time |
|---|---|---|---|---|---|---|---|---|
| tRF | 35 | GCATGGGTGGTTCAGTGGTAGAATTCTCGCCTGCC | 5' half from tRNA-Gly-GCC | 2.98 | 2.95 | 2.52 | 1.97 | decrease |
| tRF | 36 | TCCCTGGTGGTCTAGTGGTTAGGATTCTGCGCTCTC | 5' half from tRNA-Glu-CTC | 1.70 | 1.09 | 0.95 | 0.69 | decrease |
| tRF | 36 | TCCCTGGTGGTCTAGTGGCTAGGATTCGGCGCTTTC | 5' half from tRNA-Glu-TTC | 2.87 | 2.81 | 2.51 | 1.90 | decrease |
| tRF | 36 | GTTTCCGTAGTGTAGTGGTCATCACGCTCGCCTGAC | 5' half from tRNA-Val-AAC/CAC | 2.55 | 2.50 | 2.08 | 1.58 | decrease |
| tRF | 36 | ACCCTGGTGGTCTAGTGGTTAGGATTCGGCGCTCTC | 5' half from tRNA-Glu-CTC | 2.50 | 2.47 | 2.31 | 1.54 | decrease |
| tRF | 35 | GCATTGGTGGTTCAATGGTAGAATTCTCGCCTCCC | 5' half from tRNA-Gly-CCC | 2.68 | 2.67 | 2.31 | 1.75 | decrease |
| tRF | 35 | GCATTGGTAGTTCAATGGTAGAATTCTCGCCTCCG | 5' half from tRNA-Gly-CCC | 1.81 | 1.72 | 1.34 | 0.89 | decrease |
| tRF | 36 | GTTTCTGTAGTGTAGTGGTTATCACGCTCGCCTGAC | 5' half from tRNA-Val-AAC | 3.87 | 3.82 | 3.49 | 2.95 | decrease |
| tRF | 34 | CTTTCCGTAGTGTAGTGGTTATCACGTTCGCCTC | 5' half from tRNA-Val-CAC | 1.62 | 1.22 | 0.98 | 0.75 | decrease |
| tRF | 36 | TCCCTTGTGGTCTAGTGGTTAGGATTCGGCGCTCTC | 5' half from tRNA-Glu-CTC | 2.01 | 1.76 | 1.60 | 1.15 | decrease |
| tRF | 35 | GCATTGGTAGTTCAATGGTAGAATTCTCGCCTCCC | 5' half from tRNA-Gly-CCC | 3.77 | 3.71 | 3.33 | 2.91 | decrease |
| tRF | 35 | GCATTAGTAGTTCAATGGTAGAATTCTCGCCTCCC | 5' half from tRNA-Gly-CCC | 1.63 | 1.56 | 1.22 | 0.78 | decrease |
| tRF | 35 | GCGTTGGTAGTTCAATGGTAGAATTCTCGCCTCCC | 5' half from tRNA-Gly-CCC | 1.44 | 1.42 | 1.10 | 0.60 | decrease |
| tRF | 33 | GTTTCTGTAGTGTAGTGGTTATCACGCTCGCCA | 5' half fro mtRNA-Val-AAC | 2.14 | 2.13 | 1.76 | 1.31 | decrease |
| microRNA | 22 | GAGGGTTGGGTGGAGGCTCTCC | mmu-miR-296-3p | 1.87 | 1.79 | 1.66 | 1.05 | decrease |
| tRF | 35 | GCATTGGTGGTTCAGTGGTAGAATTCTCGCCTGAC | 5' half from tRNA-Gly-GCC | 1.88 | 1.75 | 1.42 | 1.07 | decrease |
| tRF | 24 | TATCGTCCTTCTGCCCAGCGATCT | tRF-1 from tRNA-His-GTG | 1.90 | 1.81 | 1.64 | 1.09 | decrease |
| tRF | 33 | GTTTCTGTAGTGTAGTGGTTATCACGCTCGCCC | 5' half fro mtRNA-Val-AAC | 2.22 | 2.17 | 1.80 | 1.44 | decrease |
| tRF | 35 | GCATTGGTAGTTCGATGGTAGAATTCTCGCCTCCC | 5' half from tRNA-Gly-CCC | 1.36 | 1.31 | 0.96 | 0.58 | decrease |
| tRF | 35 | GCATTGGTGGTTCAGTGGTAGAATTCTCGCATGCC | 5' half from tRNA-Gly-GCC | 1.62 | 1.33 | 0.98 | 0.85 | decrease |
| tRF | 35 | GCTTTGGTGGTTCAGTGGTAGAATTCTCGCCTGCC | 5' half from tRNA-Gly-GCC | 1.37 | 1.36 | 1.00 | 0.59 | decrease |
| tRF | 23 | GTTTCCGTAGTGTAGTGGTCATT | tRF-5 from tRNA-Val-AAC/CAC | 1.69 | 1.57 | 1.55 | 0.92 | decrease |
| tRF | 35 | GCATTGGTAGTTCAATGGTAGGATTCTCGCCTCCC | 5' half from tRNA-Gly-CCC | 1.45 | 1.42 | 1.03 | 0.69 | decrease |
| tRF | 35 | GCATTGGTAGTTCAATGGTAGAATTCCCGCCTCCC | 5' half from tRNA-Gly-CCC | 1.37 | 1.35 | 0.93 | 0.61 | decrease |
| tRF | 36 | GTTTCCGTAGTGTAGTGGTTATCACGCTCGCCTGAC | 5' half fro mtRNA-Val-CAC | 1.55 | 1.52 | 1.22 | 0.80 | decrease |
| tRF | 33 | GTTTCTGTAGTGTAGTGGTTATCACGCTCGCCT | 5' half fro mtRNA-Val-AAC | 3.05 | 2.98 | 2.57 | 2.29 | decrease |
| microRNA | 21 | TTAATGCTAATTGTGATAGGG | mmu-miR-155-5p | 1.40 | 1.07 | 1.03 | 0.64 | decrease |
| tRF | 20 | TCTGGACACATGTGGCTTTT | tRF-1 from tRNA-Asp-GTC | 1.48 | 1.31 | 1.30 | 0.72 | decrease |
| microRNA | 24 | TGAGAACTGAATTCCATAGGCTGC | mmu-miR-146b-5p | 1.77 | 1.47 | 1.46 | 1.03 | decrease |
| tRF | 36 | TCCCTCGTGGTCTAGTGGTTAGGATTCGGCGCTCTC | 5' half from tRNA-Glu-CTC | 1.46 | 1.17 | 1.03 | 0.72 | decrease |
| tRF | 23 | GTTTCCGTAGTGTAGTGGTCATA | tRF-5 from tRNA-Val-AAC/CAC | 1.23 | 1.04 | 1.01 | 0.50 | decrease |
| tRF | 36 | TCCCTGGTGGTCTAGTGGTTAGGATTTGGCGCTCTT | 5' half from tRNA-Glu-CTC | 1.37 | 1.06 | 1.05 | 0.64 | decrease |
| tRF | 35 | GCATTCGTGGTTCAGTGGTAGAATTCTCGCCTGCC | 5' half from tRNA-Gly-GCC | 1.72 | 1.61 | 1.21 | 0.99 | decrease |
| tRF | 35 | GCATTGGTAGTTCAATGGTGGAATTCTCGCCTCCC | 5' half from tRNA-Gly-CCC | 1.27 | 1.20 | 0.85 | 0.55 | decrease |
| tRF | 35 | TCATTGGTGGTTCAGTGGTAGAATTCTCGCCTGCC | 5' half from tRNA-Gly-GCC | 1.81 | 1.53 | 1.18 | 1.08 | decrease |
| tRF | 36 | TCCCTGGAGGTCTAGTGGTTAGGATTCGGCGCTCTC | 5' half from tRNA-Glu-CTC | 1.82 | 1.78 | 1.63 | 1.10 | decrease |
| tRF | 33 | GTTTCCGTAGTGTAGTGGTCATCACGCTCGCCG | 5' half from tRNA-Val-AAC/CAC | 2.17 | 2.05 | 1.53 | 1.45 | decrease |
| tRF | 18 | ACACTTGTCAGTTTCTTT | tRF-1 from tRNA-Val-CAC | 1.66 | 1.48 | 1.45 | 0.94 | decrease |
| tRF | 35 | GCATTGGAGGTTCAGTGGTAGAATTCTCGCCTGCC | 5' half from tRNA-Gly-GCC | 1.78 | 1.75 | 1.40 | 1.08 | decrease |
| tRF | 28 | GTTTCCGTAGTGTAGTGGTCATCACGCT | tRF-5 from tRNA-Val-AAC/CAC | 1.89 | 1.54 | 1.41 | 1.19 | decrease |
| microRNA | 25 | TGAGAACTGAATTCCATAGGCTGTT | mmu-miR-146b-5p | 1.35 | 1.09 | 1.02 | 0.65 | decrease |
| tRF | 36 | TCCCTGGTGGTCTAGTGGTTAGGATTCGGCGATCTC | 5' half from tRNA-Glu-CTC | 1.48 | 1.28 | 1.14 | 0.79 | decrease |
| tRF | 35 | GCATTGGTAGCTCAATGGTAGAATTCTCGCCTCCC | 5' half from tRNA-Gly-CCC | 1.26 | 1.19 | 0.85 | 0.56 | decrease |
| tRF | 36 | TCCCTGGTGGTCTAGTGGTTAGGATTCGGCGCTCTG | 5' half from tRNA-Glu-CTC | 1.84 | 1.83 | 1.71 | 1.15 | decrease |
| tRF | 35 | TCCTGGTGGTCTAGTGGTTAGGATTCGGCGCTCTC | 5' half from tRNA-Glu-CTC | 1.38 | 1.31 | 1.17 | 0.69 | decrease |
| microRNA | 24 | TGAGAACTGAATTCCATAGGCTGT | mmu-miR-146b-5p | 2.63 | 2.32 | 2.27 | 1.96 | decrease |
| tRF | 34 | GTTTCCGAAGTGTAGTGGTTATCACGTTCGCCTC | 5' half from tRNA-Val-CAC | 1.45 | 1.45 | 1.17 | 0.77 | decrease |
| tRF | 36 | TCCATGGTGGTCTAGTGGTTAGGATTCGGCGCTCTC | 5' half from tRNA-Glu-CTC | 2.07 | 1.70 | 1.56 | 1.40 | decrease |
| microRNA | 22 | TTAATGCTAATTGTGATAGGGG | mmu-miR-155-5p | 1.57 | 1.26 | 1.08 | 0.90 | decrease |
| tRF | 36 | TCCCTGGTGGTCTAGTGGTTATGATTCGGCGCTCTC | 5' half from tRNA-Glu-CTC | 1.31 | 1.03 | 0.86 | 0.65 | decrease |
| microRNA | 21 | ATACAGACACATGCACACACC | mmu-miR-466g-3p | 1.51 | 1.47 | 1.45 | 0.85 | decrease |
| tRF | 36 | TCCCTGGTGGTCTAGTGGTTAGGATTCGGAGCTCTC | 5' half from tRNA-Glu-CTC | 1.53 | 1.35 | 1.21 | 0.87 | decrease |
| tRF | 35 | TCCCTGGTGGTCTAGTGGTTAGGATTCTGCGCTCT | 5' half from tRNA-Glu-CTC | 1.23 | 0.76 | 0.72 | 0.57 | decrease |
| tRF | 35 | GCATTGGTGGTTCAGTGGTAGAATTCTAGCCTGCC | 5' half from tRNA-Gly-GCC | 1.50 | 1.35 | 0.95 | 0.84 | decrease |
| tRF | 36 | TCCCTGGTGGTCTAGTGGTTAGGATTAGGCGCTCTC | 5' half from tRNA-Glu-CTC | 1.67 | 1.46 | 1.29 | 1.02 | decrease |
| tRF | 36 | TACCTGGTGGTCTAGTGGTTAGGATTCGGCGCTCTC | 5' half from tRNA-Glu-CTC | 1.88 | 1.65 | 1.48 | 1.23 | decrease |
| microRNA | 23 | TGAGAACTGAATTCCATAGGCTT | mmu-miR-146b-5p | 1.33 | 1.04 | 0.97 | 0.69 | decrease |
| tRF | 36 | TCCCTGGTGGTCTAGTGGTTAGGATTTGGCGCTCTC | 5' half from tRNA-Glu-CTC | 2.35 | 1.87 | 1.72 | 1.71 | decrease |
| microRNA | 23 | TGAGAACTGAATTCCATAGGCTG | mmu-miR-146b-5p | 2.68 | 2.26 | 2.22 | 2.04 | decrease |
| tRF | 23 | GTTTCCGTAGTGTAGTGGTCATC | tRF-5 from tRNA-Val-AAC/CAC | 2.75 | 2.45 | 2.29 | 2.11 | decrease |
| tRF | 36 | TTCCTGGTGGTCTAGTGGTTAGGATTCGGCGCTCTC | 5' half from tRNA-Glu-CTC | 1.82 | 1.79 | 1.60 | 1.18 | decrease |
| microRNA | 22 | ATACACACACACATACACACTC | mmu-miR-466i-3p | 1.98 | 1.82 | 1.81 | 1.34 | decrease |
| microRNA | 23 | TTAATGCTAATTGTGATAGGGGT | mmu-miR-155-5p | 2.07 | 1.81 | 1.77 | 1.44 | decrease |
| tRF | 34 | GTTTCCGTAGTGTAGTGGTTATCACGTTCGCATC | 5' half from tRNA-Val-CAC | 1.28 | 1.12 | 0.86 | 0.65 | decrease |
| tRF | 37 | TCCCTGGTGGTCTAGTGGTTAGGATTCGGCGCTCTCT | 5' half from tRNA-Glu-CTC | 1.41 | 1.40 | 1.23 | 0.79 | decrease |
| tRF | 35 | GCATTGGTGGTTCAGTGGTAGAATTCTTGCCTGCC | 5' half from tRNA-Gly-GCC | 1.85 | 1.63 | 1.30 | 1.24 | decrease |
| tRF | 36 | TCCCTGGTGGTCTAGTGGTTAGGATTCGCCGCTCTC | 5' half from tRNA-Glu-CTC | 1.35 | 1.24 | 1.02 | 0.74 | decrease |
| tRF | 36 | TCCTTGGTGGTCTAGTGGTTAGGATTCGGCGCTCTC | 5' half from tRNA-Glu-CTC | 1.68 | 1.64 | 1.48 | 1.07 | decrease |
| microRNA | 23 | GGCAGAGGAGGGCTGTTCTTCCC | mmu-miR-298-5p | 2.18 | 2.07 | 1.99 | 1.57 | decrease |
| tRF | 36 | GTTTCCGTAGTGTAGTGGTCATCACGCTCGCCTCAT | 5' half from tRNA-Val-CAC | 1.20 | 1.14 | 0.99 | 0.60 | decrease |
| tRF | 39 | ACCGCCGCGGCCCGGGTTCGATTCCCGGTCAGGGAACCA | 3' half from tRNA-Glu-CTC | 2.13 | 1.99 | 1.98 | 1.53 | decrease |
| tRF | 36 | GTTTCCGTAGTGTAGTGGTTATCACATTCGCCTGAC | 5' half from tRNA-Val-AAC | 2.31 | 2.29 | 1.94 | 1.72 | decrease |
| tRF | 36 | TCCCTGGTGGTCTAGTGGCTAGGATTCGGCGCTCTC | 5' half from tRNA-Glu-CTC | 2.03 | 2.01 | 1.85 | 1.45 | decrease |
| tRF | 15 | TCTAGGATTTCTTTT | tRF-1 from tRNA-Ser-AGA | 1.40 | 1.05 | 1.04 | 0.82 | decrease |
| tRF | 36 | GCCCTGGTGGTCTAGTGGTTAGGATTCGGCGCTCTC | 5' half from tRNA-Glu-CTC | 1.88 | 1.71 | 1.54 | 1.30 | decrease |
| tRF | 32 | GTTTCTGTAGTGTAGTGGTTATCACGCTCGCC | 5' half from tRNA-Val-AAC | 2.34 | 2.26 | 1.85 | 1.77 | decrease |
| tRF | 36 | GTTTCCGTAGTGTAGTGGTTATCACGTTCGCCTGAC | 5' half from tRNA-Val-AAC/CAC | 3.27 | 3.23 | 2.93 | 2.70 | decrease |

| | | | | | | | | |
|---|---|---|---|---|---|---|---|---|
| tRF | 33 | GTTTCAGTAGTGTAGTGGTTATCACGCTCGCCT | 5' half from tRNA-Val-AAC | 1.18 | 1.09 | 0.77 | 0.61 | decrease |
| tRF | 36 | TCCCTGGTGGTCTAGTGGTTAGGATTCAGCGCTCTC | 5' half from tRNA-Glu-CTC | 1.84 | 1.83 | 1.65 | 1.27 | decrease |
| microRNA | 24 | TTTTGCAGTATGTTCCTGAATACA | mmu-miR-450b-5p | 1.78 | 1.56 | 1.56 | 1.21 | decrease |
| tRF | 36 | GTTTCCGTAGTGTAGTGGTCATCACGCTCGCCTCAC | 5' half from tRNA-Val-CAC | 1.95 | 1.73 | 1.44 | 1.38 | decrease |
| tRF | 34 | GTTTCCGTAGTGTAGTGGTTATCACGCTCGCCTC | 5' half from tRNA-Val-CAC | 2.91 | 2.87 | 2.50 | 2.36 | decrease |
| tRF | 36 | TCCCTGGTGGTTTAGTGGTTAGGATTCGGCGCTCTC | 5' half from tRNA-Glu-CTC | 1.59 | 1.51 | 1.35 | 1.03 | decrease |
| tRF | 36 | TCCCGGGTGGTCTAGTGGTTAGGATTCGGCGCTCTC | 5' half from tRNA-Glu-CTC | 1.25 | 1.19 | 1.06 | 0.70 | decrease |
| tRF | 36 | TCCCTGGTGGTCTAGTGGTTAGGATTCGGCGCTTTC | 5' half from tRNA-Glu-CTC | 1.66 | 1.50 | 1.37 | 1.11 | decrease |
| tRF | 36 | TCCCTGGTGGTCTGGTGGTTAGGATTCGGCGCTCTC | 5' half from tRNA-Glu-CTC | 2.05 | 2.05 | 1.89 | 1.50 | decrease |
| tRF | 36 | GTTTCCGTAGTGTAGTGGTTATCACGTTCGCCTCAT | 5' half from tRNA-Val-CAC | 2.23 | 2.19 | 2.14 | 1.68 | decrease |
| tRF | 36 | TCCCTGGTGGTCTAGTGGTTAAGATTCGGCGCTCTC | 5' half from tRNA-Glu-CTC | 1.55 | 1.53 | 1.37 | 1.00 | decrease |
| tRF | 35 | GCATTGGTGGTTCAGTGGTAGAATTCTCGCCTGTC | 5' half from tRNA-Gly-GCC | 1.77 | 1.74 | 1.38 | 1.22 | decrease |
| tRF | 36 | TCCCTGGTGGTCTAGTGGTTAGGATTCGGTGCTCTC | 5' half from tRNA-Glu-CTC | 1.88 | 1.87 | 1.70 | 1.33 | decrease |
| tRF | 36 | TCCCTGGTGGTATAGTGGTTAGGATTCGGCGCTCTC | 5' half from tRNA-Glu-CTC | 1.48 | 1.38 | 1.26 | 0.93 | decrease |
| tRF | 36 | TCCCTGGTGGTCTAGTGGTTAGGATTCGACGCTCTC | 5' half from tRNA-Glu-CTC | 1.61 | 1.57 | 1.44 | 1.06 | decrease |
| tRF | 34 | GTTTCAGTAGTGTAGTGGTTATCACGTTCGCCTC | 5' half from tRNA-Val-CAC | 1.96 | 1.68 | 1.44 | 1.41 | decrease |
| tRF | 35 | GCATTGGTGGTTCAGTGGTAGAATTCCCGCCTGCC | 5' half from tRNA-Gly-GCC | 2.10 | 2.10 | 1.73 | 1.56 | decrease |
| microRNA | 24 | TGGAAGACTAGTGATTTTGTTGTG | mmu-miR-7a-1/2-5p | 1.79 | 1.55 | 1.48 | 1.26 | decrease |
| microRNA | 22 | TGAGAACTGAATTCCATAGGCA | mmu-miR-146b-5p | 1.46 | 1.14 | 1.10 | 0.93 | decrease |
| microRNA | 24 | TTAATGCTAATTGTGATAGGGGTT | mmu-miR-155-5p | 1.93 | 1.65 | 1.62 | 1.40 | decrease |
| microRNA | 25 | TTAATGCTAATTGTGATAGGGGTTT | mmu-miR-155-5p | 1.37 | 1.12 | 1.06 | 0.84 | decrease |
| tRF | 35 | GCATTGGTGGTTCAGTGGTAGAATTCTCGTCTGCC | 5' half from tRNA-Gly-GCC | 1.56 | 1.54 | 1.20 | 1.04 | decrease |
| microRNA | 24 | CAAAGTGCTTACAGTGCAGGTAGA | mmu-miR-17-5p | 2.28 | 2.06 | 1.99 | 1.75 | decrease |
| tRF | 33 | GTTTCTCTAGTGTAGTGGTTATCACGCTCGCCT | 5' half from tRNA-Val-AAC | 1.57 | 1.53 | 1.18 | 1.04 | decrease |
| tRF | 33 | GTTTCCGTAGTGTAGTGGTCGTCACGCTCGCCT | 5' half from tRNA-Val-AAC/CAC | 1.16 | 1.15 | 0.86 | 0.64 | decrease |
| microRNA | 22 | GCCTGCTGGGGTGGAACCTGGG | mmu-miR-370-3p | 1.55 | 1.30 | 1.24 | 1.02 | decrease |
| tRF | 23 | TATCGTCCTTCTGCCCAGCGATC | tRF-1 from tRNA-His-GTG | 1.77 | 1.69 | 1.60 | 1.24 | decrease |
| tRF | 30 | TCCCTGGTGGTCTAGTGGCTAGGATTCGGC | 5' half from tRNA-Glu-TTC | 1.92 | 1.57 | 1.53 | 1.40 | decrease |
| tRF | 36 | TCCCTGGTGGTCTAGTGGTTAGGATTCGGCGCTCGC | 5' half from tRNA-Glu-CTC | 1.22 | 1.13 | 0.98 | 0.70 | decrease |
| tRF | 35 | GCATTGGTGGTTCAGTGGTAAAATTCTCGCCTGCC | 5' half from tRNA-Gly-GCC | 1.47 | 1.41 | 1.10 | 0.95 | decrease |
| tRF | 16 | GTTTCCGTAGTGTAGT | tRF-5 from tRNA-Val-AAC/CAC | 1.69 | 1.41 | 1.31 | 1.18 | decrease |
| tRF | 35 | TCCCTTGTGGTCTAGTGGTTAGGATTCGGCGCTCT | 5' half from tRNA-Glu-CTC | 1.52 | 1.32 | 1.28 | 1.01 | decrease |
| tRF | 36 | GGTTCCATAGTGTAGCGGTTATCACGTCTGCTTTAC | 5' half from tRNA-Val-TAC | 1.47 | 1.27 | 1.06 | 0.96 | decrease |
| microRNA | 21 | TGAGAACTGAATTCCATAGGC | mmu-miR-146b-5p | 1.53 | 1.20 | 1.12 | 1.01 | decrease |
| tRF | 17 | GTTTCCGTAGTGTAGTG | tRF-5 from tRNA-Val-AAC/CAC | 2.14 | 1.90 | 1.78 | 1.63 | decrease |
| tRF | 34 | GTTTCCGTAGTGTAGTGGTTATCACACTCGCCTC | 5' half fro mtRNA-Val-CAC | 1.37 | 1.34 | 1.01 | 0.86 | decrease |
| microRNA | 22 | CTCACCTGGAGCATGTTTTCTT | mmu-miR-1983-3p | 1.43 | 1.41 | 1.37 | 0.92 | decrease |
| tRF | 35 | TTGTGGCAACAATGGTACGGCAAGGGCGCCTTTTT | tRF-1 from tRNA-His-GTG | 1.28 | 1.11 | 1.09 | 0.78 | decrease |
| microRNA | 21 | CCTGAACTAGGGGTCTGGAGA | mmu-miR-345-3p | 1.34 | 1.12 | 1.12 | 0.84 | decrease |
| microRNA | 23 | TAATACTGTCTGGTAATGCCGTT | mmu-miR-429-3p | 0.93 | 0.98 | 1.27 | 1.43 | increase |
| microRNA | 20 | TTTGAACCATCACTCGACTC | mmu-miR-434-3p | 1.55 | 1.61 | 1.86 | 2.05 | increase |
| microRNA | 23 | CGGATCCGTCTGAGCTTGGCTAT | mmu-miR-127-3p | 0.68 | 0.69 | 1.04 | 1.19 | increase |
| tRF | 25 | TTGGTGGTTCAGTGGTAGAATTCTC | i-tRF from tRNA-Gly-GCC | 0.66 | 0.75 | 0.86 | 1.18 | increase |
| microRNA | 22 | TAATACTGCCGGGTAATGATGA | mmu-miR-200c-3p | 1.50 | 1.58 | 1.94 | 2.02 | increase |
| microRNA | 23 | TGGTAGACTATGGAACGTAGGAA | mmu-miR-379-5p | 0.85 | 0.91 | 1.28 | 1.37 | increase |
| microRNA | 23 | TCGGATCCGTCTGAGCTTGGCAT | mmu-miR-127-3p | 1.45 | 1.56 | 1.92 | 1.98 | increase |
| microRNA | 20 | TAATACTGCCGGGTAATGAC | mmu-miR-200c-3p | 0.74 | 0.79 | 1.11 | 1.29 | increase |
| microRNA | 23 | TGTAACAGCAACTCCATGTGGAT | mmu-miR-194-1/2-5p | 1.06 | 1.18 | 1.60 | 1.63 | increase |
| microRNA | 25 | TCCCTGAGGAGCCCTTTGAGCCTTT | mmu-miR-351-5p | 1.64 | 1.65 | 1.92 | 2.21 | increase |
| microRNA | 19 | GAAGTTGTTCGTGGTGGAT | mmu-miR-382-5p | 0.83 | 0.85 | 1.07 | 1.41 | increase |
| microRNA | 24 | CCCAGTGTTCAGACTACCTGTTCT | mmu-miR-199a-1/2-5p | 1.26 | 1.27 | 1.49 | 1.84 | increase |
| tRF | 34 | GCCGTGATCGTATAGTGGTTAGTACTCTGCGTTG | 5' half from tRNA-His-GTG | 1.68 | 1.72 | 1.75 | 2.26 | increase |
| microRNA | 21 | TCCCTGAGGAGCCCTTTGAGA | mmu-miR-351-5p | 1.21 | 1.27 | 1.47 | 1.81 | increase |
| microRNA | 24 | TGACCTATGAATTGACAGCCAGTT | mmu-miR-192-5p | 0.63 | 0.74 | 1.04 | 1.23 | increase |
| microRNA | 21 | TCCCTGAGGAGCCCTTTGAGT | mmu-miR-351-5p | 1.95 | 2.07 | 2.31 | 2.57 | increase |
| microRNA | 23 | TGTAACAGCAACTCCATGTGGAC | mmu-miR-194-2-5p | 1.23 | 1.38 | 1.77 | 1.90 | increase |
| microRNA | 23 | CGGATCCGTCTGAGCTTGGCTTT | mmu-miR-127-3p | 0.78 | 0.84 | 1.41 | 1.47 | increase |
| microRNA | 21 | TGTCACTCGGCTCGGCCCACT | mmu-miR-668-3p | 0.65 | 0.71 | 1.00 | 1.36 | increase |
| microRNA | 23 | GTAAAGGCTGGGCTTAGACGTGT | mmu-miR-1981-5p | 0.87 | 1.00 | 1.47 | 1.59 | increase |
| microRNA | 22 | TCCCTGAGGAGCCCTTTGAGTT | mmu-miR-351-5p | 0.86 | 0.96 | 1.21 | 1.58 | increase |
| microRNA | 23 | CTGACCTATGAATTGACAGCCTT | mmu-miR-192-5p | 0.92 | 0.99 | 1.42 | 1.64 | increase |
| microRNA | 24 | CTGACCTATGAATTGACAGCCATC | mmu-miR-192-5p | 1.05 | 1.15 | 1.63 | 1.78 | increase |
| microRNA | 21 | TTTTGCGATGTGTTCCTAATT | mmu-miR-450a-1/2-5p | 0.73 | 0.75 | 0.96 | 1.47 | increase |
| microRNA | 20 | TCTCTGGGCCTGTGTCTTAG | mmu-miR-330-5p | 0.97 | 1.06 | 1.55 | 1.77 | increase |
| microRNA | 24 | TGTAACAGCAACTCCATGTGGACT | mmu-miR-194-2-5p | 0.80 | 0.94 | 1.39 | 1.64 | increase |
| microRNA | 23 | TGACCTATGAATTGACAGCCATT | mmu-miR-192-5p | 0.96 | 1.07 | 1.51 | 1.81 | increase |
| microRNA | 22 | ATGACCTATGATTTGACAGACC | mmu-miR-215-5p | 2.23 | 2.31 | 2.79 | 3.10 | increase |
| microRNA | 24 | CTGACCTATGAATTGACAGCCATT | mmu-miR-192-5p | 1.91 | 1.95 | 2.42 | 2.79 | increase |
| microRNA | 22 | ATGACCTATGATTTGACAGACA | mmu-miR-215-5p | 1.40 | 1.58 | 2.17 | 2.29 | increase |
| microRNA | 22 | TAATACTGCCGGGTAATGATGT | mmu-miR-200c-3p | 0.89 | 0.91 | 1.27 | 1.78 | increase |
| microRNA | 21 | TAATACTGCCGGGTAATGATA | mmu-miR-200c-3p | 0.94 | 1.05 | 1.55 | 1.98 | increase |
| microRNA | 21 | ATGACCTATGATTTGACAGAT | mmu-miR-215-5p | 0.81 | 0.94 | 1.64 | 1.88 | increase |
| microRNA | 17 | ATGACCTATGATTTGAC | mmu-miR-215-5p | 0.89 | 0.96 | 1.44 | 2.01 | increase |
| microRNA | 20 | TAACACTGTCTGGTAAAGAT | mmu-miR-141-3p | 0.60 | 0.67 | 1.12 | 1.73 | increase |
| microRNA | 21 | TAACACTGTCTGGTAAAGATG | mmu-miR-141-3p | 1.12 | 1.22 | 1.75 | 2.34 | increase |
| microRNA | 20 | ATGACCTATGATTTGACAGA | mmu-miR-215-5p | 0.75 | 0.84 | 1.47 | 2.23 | increase |

**Table S3. MIA-responsive placental/decidual microRNA and tRF sequences from DESeq2 (padj<0.05, |log2foldchange|>0.5).**

| Sequence | Annotation | baseMean | log2FoldChange (MIA/control) | lfcSE | pvalue | padj |
|---|---|---|---|---|---|---|
| GTTTCCGTAGTGTAGTGGTTATCACGTTCGCCTA | 5' half from tRNA-Val-AAC | 4843.63 | -0.88 | 0.21 | 1.50E-06 | 2.44E-04 |
| GTTTCCCTAGTGTAGTGGTTATCACGTTCGCCTC | 5' half from tRNA-Val-CAC | 129.06 | -0.85 | 0.20 | 7.96E-07 | 1.72E-04 |
| GTTTCCGTAGTGTAGTGGTTATCACATTCGCCTA | 5' half from tRNA-Val-AAC | 85.14 | -0.83 | 0.31 | 2.47E-04 | 1.04E-02 |
| GGTAGCGTGGCCGAGTGGTCTAAGGCGCTGGATTT | 5' half from tRNA-Leu-TAG | 139.49 | -0.81 | 0.23 | 1.98E-05 | 1.90E-03 |
| GTCAGGATGGCCGAGCGGTCTAAGGCGCTGCGTTC | 5' half from tRNA-Leu-CAG | 708.48 | -0.76 | 0.24 | 5.81E-05 | 3.98E-03 |
| AGCAGAGTGGCGCAGCGGAAGCGTGCTGGGCCC | 5' half from tRNA-iMet-CAT | 108.68 | -0.75 | 0.17 | 9.16E-07 | 1.88E-04 |
| CATTGGTGGTTCAGTGGTAGAATTCTCGCCT | 5' half from tRNA-Gly-CCC/GCC | 26.24 | -0.72 | 0.36 | 1.33E-03 | 3.10E-02 |
| TTGGGAACATTTTGCATGCAC | mmu-miR-450b-3p | 34.79 | -0.72 | 0.22 | 5.53E-05 | 3.85E-03 |
| GTTTCCGTAGTGTAGTGGTCATCACGCTCGCCTA | 5' half from tRNA-Val-AAC | 636.03 | -0.71 | 0.19 | 9.21E-06 | 1.11E-03 |
| GCATGGGTGGTTCAGTGGTAGAATTCTCGCCTGCC | 5' half fro tRNA-Gly-GCC | 5520.43 | -0.70 | 0.14 | 7.18E-08 | 2.58E-05 |
| GCCCGGATGATCCTCAGTGGTCTGGGGTGCAGGCTTC | 5' half from tRNA-SeC-TCA | 462.55 | -0.70 | 0.14 | 2.91E-08 | 1.14E-05 |
| TCTAGGATTTCTTTT | tRF-1 from tRNA-Ser-AGA | 136.99 | -0.69 | 0.22 | 8.08E-05 | 5.13E-03 |
| TCCCATATGGTCTAGCGGTTAGGATTCCTGGTTTTC | 5' half from tRNA-Glu-TTC | 4218.12 | -0.69 | 0.12 | 1.72E-09 | 1.24E-06 |
| TCCCACATGGTCTAGCGGTTAGGATTCCTGGTTTT | 5' half from tRNA-Glu-TTC | 1279.64 | -0.67 | 0.19 | 1.61E-05 | 1.66E-03 |
| GCATTGGTAGTTCAATGGTAGAATTCTCGCCT | 5' half from tRNA-Gly-CCC | 4861.11 | -0.65 | 0.21 | 7.13E-05 | 4.67E-03 |
| GTTTCCGTAGTGTAGTGGTTATCACGTTCGCCTC | 5' half from tRNA-Val-CAC | 72499.42 | -0.65 | 0.20 | 6.47E-05 | 4.30E-03 |
| TCCCATATGGTCTAGCGGTTAGGATTCCTGGTTTT | 5' half from tRNA-Glu-TTC | 480.75 | -0.65 | 0.19 | 3.99E-05 | 3.19E-03 |
| TCCCACATGGTCTAGCGGTTAGGATTCCTGGTTTTC | 5' half from tRNA-Glu-TTC | 13409.25 | -0.64 | 0.12 | 1.08E-08 | 5.82E-06 |
| GCATTGGTAGTTCAATGGTAGAATTCTCGCCTCCC | 5' half from tRNA-Gly-CCC | 34745.90 | -0.63 | 0.16 | 3.92E-06 | 5.46E-04 |
| TCCCACATGGTCTAGC | tRF-5 from tRNA-Glu-TTC | 33.17 | -0.62 | 0.27 | 9.63E-04 | 2.60E-02 |
| GGGGTATAGCTCAGTGGTAGAGCATTTGACTGC | 5' half from tRNA-Cys-GCA | 206.47 | -0.62 | 0.16 | 5.26E-06 | 7.09E-04 |
| GCATTGGTAGTTCAATGGTAGAATTCTCGCCTCC | 5' half from tRNA-Gly-CCC | 10918.18 | -0.61 | 0.14 | 1.11E-06 | 2.18E-04 |
| TCTCCCAACCCTTGTACCAGT | mmu-miR-150-5p | 65.69 | -0.61 | 0.18 | 3.21E-05 | 2.71E-03 |
| TCCTCGTTAGTATAGTGGTTAGTATCCCCGCCTGTC | 5' half from tRNA-Asp-GTC | 1399.48 | -0.61 | 0.13 | 1.71E-07 | 5.26E-05 |
| GTTTCCGTAGTGTAGTGGTTATCACGTTCGCCTCA | 5' half from tRNA-Val-CAC | 610.28 | -0.61 | 0.22 | 2.45E-04 | 1.04E-02 |
| GCATTGGTAGTTCAATGGTAGAATTCTCGCCTC | 5' half from tRNA-Gly-CCC | 9645.17 | -0.61 | 0.17 | 1.42E-05 | 1.54E-03 |
| GGGGTATAGCTCAGTGGTAGAGCATTTGACTG | 5' half from tRNA-Cys-GCA | 38.94 | -0.61 | 0.25 | 6.36E-04 | 1.96E-02 |
| TCCTCGTTAGTATAGTGGTGAGTATCCCCGCCTGTCA | 5' half from tRNA-Asp-GTC | 23.66 | -0.59 | 0.29 | 1.56E-03 | 3.46E-02 |
| TCCTCGTTAGTATAGTGGTTAGTATCCCCGCCTGT | 5' half from tRNA-Asp-GTC | 154.78 | -0.59 | 0.21 | 2.67E-04 | 1.09E-02 |
| GCATTGGTGGTTCAGTGGTAGAATTCTCGCCTGCC | 5' half from tRNA-Gly-GCC | 189499.26 | -0.58 | 0.15 | 1.54E-05 | 1.62E-03 |
| GCATTGGTGGTTCAGTGGTAGAATTCTCGCCT | 5' half from tRNA-Gly-CCC/GCC | 213256.96 | -0.58 | 0.17 | 4.81E-05 | 3.58E-03 |
| AGTCGGTAGAGCATCAGA | i-tRF from tRNA-Lys-TTT | 39.06 | -0.56 | 0.19 | 1.45E-04 | 7.13E-03 |
| GTTTCCGTAGTGTAGTGGTCATCACGCTCGCCTCA | 5' half from tRNA-Val-CAC | 55.39 | -0.55 | 0.23 | 8.56E-04 | 2.42E-02 |
| ATGGGTGGTTCAGTGGTAGAATTCTCGCCTGC | 5' half from tRNA-Gly-GCC | 32.27 | -0.55 | 0.24 | 1.08E-03 | 2.73E-02 |
| AAAGGACTATTTTTTT | tRF-1 from tRNA-Arg-CCT | 80.95 | -0.55 | 0.26 | 1.60E-03 | 3.46E-02 |
| GGTTCCATAGTGTAGCGGTTATCACGTCTGCTTTAC | 5' half from tRNA-Val-TAC | 173.91 | -0.54 | 0.14 | 6.10E-06 | 7.74E-04 |
| GCATTGGTGGTTCAATGGTAGAATTCTCGCCT | 5' half from tRNA-Gly-CCC | 420.01 | -0.54 | 0.20 | 3.12E-04 | 1.24E-02 |
| CATTATTACTTTTGGTACGT | mmu-miR-126a-5p | 39.35 | -0.54 | 0.24 | 1.20E-03 | 2.92E-02 |
| GCCCGGCTAGCTCAGTCGGTAGAGCATGAGACTCTTAAT | 5' half from tRNA-Lys-CTT | 45.12 | -0.54 | 0.23 | 9.31E-04 | 2.58E-02 |
| TCCTCGTTAGTATAGTGGTGAGTATCCCCGCCTGTC | 5' half from tRNA-Asp-GTC | 9773.47 | -0.52 | 0.13 | 3.43E-06 | 5.11E-04 |
| GCCCGGCTAGCTCAGTCGGTAGAGCATGAGACTC | 5' half from tRNA-Lys-CTT | 26955.19 | -0.51 | 0.20 | 6.20E-04 | 1.96E-02 |
| GCCCGGCTAGCTCAGTCGGTAGAGCATGGGACTC | 5' half from tRNA-Lys-CTT | 2177.40 | -0.50 | 0.22 | 1.07E-03 | 2.72E-02 |
| GCATTGGTGGTTCAGTGGTAGAATTCTCGCCTC | 5' half from tRNA-Gly-CCC | 83.68 | -0.50 | 0.24 | 1.70E-03 | 3.65E-02 |
| TCCCCGGCATCTCCACCA | tRF-3 from tRNA-Ala-CGC/TGC | 2487.97 | 0.50 | 0.08 | 1.79E-10 | 1.94E-07 |
| GCTCTGACTAGGTTGCACTACT | mmu-miR-301b-5p | 37.26 | 0.50 | 0.25 | 1.89E-03 | 3.94E-02 |
| CCAATATTGGCTGTGCTGCTCC | mmu-miR-195a-3p | 92.83 | 0.51 | 0.21 | 7.46E-04 | 2.19E-02 |
| ACCACAGGGTAGAACC | mmu-miR-140-3p | 66.65 | 0.51 | 0.25 | 1.82E-03 | 3.86E-02 |
| TGCCTGTCTACACTTGCTGTGC | mmu-miR-214-5p | 179.86 | 0.51 | 0.14 | 1.95E-05 | 1.90E-03 |
| TGGAGGCGTGGGTTCG | i-tRF from tRNA-Leu-CAA/CAG | 24.79 | 0.51 | 0.28 | 2.54E-03 | 4.70E-02 |
| ATCGATGTGGTGCTCC | mmu-miR-5099-3p | 116.22 | 0.52 | 0.20 | 5.79E-04 | 1.89E-02 |
| ACCGGGCGGGAACACC | tRF-3 from tRNA-Val-CAC/AAC | 53.78 | 0.52 | 0.20 | 4.72E-04 | 1.62E-02 |
| AAGCTCGGTCTGAGGCCCCTCA | mmu-miR-423-3p | 56.07 | 0.52 | 0.17 | 1.38E-04 | 7.02E-03 |
| CGTATGGAGGCGTGGGT | i-tRF from tRNA-Leu-CAA | 26.21 | 0.54 | 0.24 | 1.18E-03 | 2.90E-02 |
| GTGCCTACTGAGCTGAAA | miR-24-2-5p | 31.16 | 0.54 | 0.25 | 1.22E-03 | 2.96E-02 |
| TGGTTAGGATTCGGCGCTCTCACC | i-tRF from tRNA-Glu-CTC | 51.58 | 0.55 | 0.21 | 4.69E-04 | 1.62E-02 |
| CCACTTCTGACACCA | mmu-miR-3963-5p | 51.46 | 0.57 | 0.18 | 7.44E-05 | 4.80E-03 |
| CCGACTTCTGGGCTCCGGCTTTT | mmu-miR-1964-3p | 43.19 | 0.57 | 0.24 | 7.90E-04 | 2.27E-02 |
| ATCTCGGTGGAACCTCC | tRF-3 from tRNA-Gln-CTG | 60.55 | 0.57 | 0.17 | 5.23E-05 | 3.77E-03 |
| CATCAAAGTGGAGGCCCTCT | mmu-miR-291a-5p | 244.04 | 0.58 | 0.12 | 8.30E-08 | 2.76E-05 |
| ATCCCACTTCTGACACCA | mmu-miR-3963-5p | 2450.22 | 0.59 | 0.17 | 4.33E-05 | 3.34E-03 |
| ATCCTGCCGACTACGCCA | tRF-3 from tRNA-Ser-AGA/TGA | 253.23 | 0.59 | 0.15 | 3.92E-06 | 5.46E-04 |
| TGGAGGCGTGGGTTC | i-tRF from tRNA-Leu-CAA/CAG | 38.61 | 0.59 | 0.22 | 3.20E-04 | 1.26E-02 |

| | | | | | | |
|---|---|---|---|---|---|---|
| CCGTGATCGTATAGTGGTTAGTACTCTGC | 5' half from tRNA-His-GTG | 192.15 | 0.61 | 0.20 | 8.77E-05 | 5.49E-03 |
| GGTCCCATGGTGTAATGGTTAGCACTCTG | tRF-5 from tRNA-Gln-TTG | 35.57 | 0.62 | 0.21 | 1.28E-04 | 6.64E-03 |
| ATCTCGGTGGGACCTCC | tRF-3 from tRNA-Gln-CTG/TTG | 38.56 | 0.62 | 0.19 | 4.28E-05 | 3.34E-03 |
| GTAAAGGCTGGGCTTAGACGTGG | mmu-miR-1981-5p | 516.06 | 0.62 | 0.18 | 2.87E-05 | 2.48E-03 |
| ATCACGTCGGGGTCACCA | tRF-3 from tRNA-Trp-CCA | 67.20 | 0.63 | 0.22 | 1.84E-04 | 8.63E-03 |
| ATCTCGCTGGGGCCTCCA | tRF-3 from tRNA-Thr-TGT | 177.49 | 0.64 | 0.16 | 5.46E-06 | 7.14E-04 |
| TCTCGCTGGGGCCTCC | tRF-3 from tRNA-Thr-TGT | 164.09 | 0.65 | 0.15 | 5.19E-07 | 1.32E-04 |
| CGTGATCGTATAGTGGTTAGTACTCTGC | 5' half from tRNA-His-GTG | 92.27 | 0.66 | 0.25 | 3.24E-04 | 1.26E-02 |
| GATCAAAGTGGAGGCCCTCA | mmu-miR-291b-5p | 26.10 | 0.67 | 0.23 | 1.42E-04 | 7.13E-03 |
| GATCAAAGTGGAGGCCCTCC | mmu-miR-291b-5p | 57.40 | 0.68 | 0.16 | 1.68E-06 | 2.60E-04 |
| GCCGTGATCGTATAGTGGTTAGTACTCTGC | 5' half from tRNA-His-GTG | 579.16 | 0.69 | 0.27 | 3.95E-04 | 1.47E-02 |
| ATCCGGCTCGAAGGACCA | tRF-3 from tRNA-Tyr-GTA | 186.14 | 0.75 | 0.12 | 2.58E-11 | 3.72E-08 |
| CCCAGCGGTGCCTCCA | mmu-miR-5100-3p | 86.19 | 0.76 | 0.15 | 1.43E-08 | 6.86E-06 |
| TTCCGGCTCGAAGGACCA | tRF-3 from tRNA-Tyr-GTA | 543.26 | 0.76 | 0.11 | 4.51E-13 | 9.74E-10 |
| ATCTCGGTGGGACCTCCA | tRF-3 from tRNA-Gln-CTG | 569.54 | 0.76 | 0.14 | 6.00E-09 | 3.70E-06 |
| TCGATCAAAGTGGAGGCCCTCT | mmu-miR-291b-5p | 65.84 | 0.77 | 0.18 | 1.27E-06 | 2.22E-04 |
| GATTTCTGCCCAGTGCTCTG | mmu-miR-6240-3p | 39.19 | 0.79 | 0.22 | 1.71E-05 | 1.72E-03 |
| ATCTCGGTGGAACCTCCA | tRF-3 from tRNA-Gln-CTG | 987.66 | 0.83 | 0.14 | 3.82E-10 | 3.30E-07 |
| ATCCCAGCGGTGCCTCCA | mmu-miR-5100-3p | 93.04 | 0.83 | 0.20 | 1.29E-06 | 2.22E-04 |
| TCCGAGCCTGGGTCTCCCT | mmu-miR-615-3p | 38.27 | 0.95 | 0.21 | 4.22E-07 | 1.22E-04 |
| CTCCGGCTCGAAGGACCA | tRF-3 from tRNA-Tyr-GTA | 100.33 | 0.96 | 0.19 | 1.69E-08 | 7.31E-06 |

**Table S4. Probe/primer sequences**

| Target | Probe/primer sequence |
|---|---|
| ***Northern blots*** | |
| 5’ tRNA-GluCTC: | 5'-BIO-CCTAACCACTAGACCACCAGGGA-3' |
| 5’ tRNA-GlyGCC: | 5'-BIO-TCTACCACTGAACCACCCATGC-3' |
| 5' tRNA-GlyC/GCC: | 5'-BIO-GAATTCTACCACTGAACCACCAATGC-3' |
| 5' tRNA-LysCTT: | 5'-BIO-CTCTACCGACTGAGCTAGCCGGGC-3' |
| 5' tRNA-ValA/CAC: | 5'-BIO-BIO-GATAACCACTACACTACGGAAAC-3' |
| 5’ tRNA-HisGTG: | 5'-BIO-CAGAGTACTAACCACTATACGATCACGGC-3' |
| 5' tRNA-AspGTC: | 5'-BIO-GGATACTCACCACTATACTAACGAGGA-3' |
| 3' tRNA-AspGTC: | 5'-BIO-GTCGGGGAATCGAACCCCGGTC-3' |
| 3' tRNA-GluC/TTC: | 5'-BIO-GTTCCCTGACCGGGAATCGAAC-3' |
| 3' tRNA-AlaC/TGC: | 5'-BIO-TGGTGGAGATGCCGGGGA-3' |
| 3' tRNA-AlaA/TGC: | 5'-BIO-TGGTGGAGGTGCCGGGGA-3' |
| 3' tRNA-GlnCTG: | 5'-BIO-TGGAGGTTCCACCGAGAT-3' |
| 3' tRNA-ValA/CAC: | 5'-BIO-TGGTGTTTCCGCCCGGT-3' |
| 3' tRNA-LeuA/TAG: | 5'-BIO-TGGTGGCAGCGGTGGGAT-3' |
| 3' tRNA-CysGCA: | 5'-BIO-TGGAGGGGGCACCCGGA-3' |
| ***qRT-PCR*** | |
| NEBNext 3' SR adaptor | 5'-rAppAGATCGGAAGAGCACACGTCT-NH2-3' |
| NEBNext 5' SR adaptor | 5'-rGrUrUrCrArGrArGrUrUrCrUrArCrArGrUrCrCrGrArCrGrArUrC-3' |
| NEBNext SR RT Primer | 5'-AGACGTGTGCTCTTCCGATCT-3' |
| mmu-miR-26a-5p | Forwad: 5'-CGATCTTCAAGTAATCCAGGA -3' |
| | Reverse: 5'-AGACGTGTGCTCTTCCGATCT-3' |
| mmu-miR-21a-5p | Forward: 5'-CGATCTAGCTTATCAGACTGA-3' |
| | Reverse: 5'-AGACGTGTGCTCTTCCGATCT-3' |
| tRF3a-Ala | Forward: 5'-CGATCTCCCCGGCATCTCCA-3' |
| | Reverse: 5'-AGACGTGTGCTCTTCCGATCT-3' |
| tRF3a-Tyr | Forwad: 5'-CGATCTTCCGGCTCGAAGGA-3' |
| | Reverse: 5'-AGACGTGTGCTCTTCCGATCT-3' |
| 5'tRH-GluTTC | Forward: 5'-CGATCTCCCTGGTGGTCTA-3' |
| | Reverse: 5'-CGATCTGAAAGCGCCGAATC-3' |
| 5'tRH-GlyCCC | Forward: 5'-GACGATCGCATTGGTAGTTCA-3' |
| | Reverse: 5'-CGATCTGGGAGGCGAGAA-3' |
| 5'tRH-AspGTC | Forward: 5'-CGATCTCCTCGTTAGTATAGTGGTGA-3' |
| | Reverse: 5'-CGATCTGACAGGCGGGGATAC-3' |
| 5'tRH-GluTTC-2 | Forward: 5'-CGATCTCCCATATGGTCTAGC-3' |
| | Reverse: 5'-CGATCTGAAAACCAGGAATCCTA-3' |
